# Bacterial Argonaute nucleases reveal different modes of DNA targeting *in vitro* and *in vivo*

**DOI:** 10.1101/2022.09.09.507302

**Authors:** Lidiya Lisitskaya, Ekaterina Kropocheva, Aleksei Agapov, Maria Prostova, Vladimir Panteleev, Denis Yudin, Sergey Ryazansky, Anton Kuzmenko, Alexei A. Aravin, Daria Esyunina, Andrey Kulbachinskiy

## Abstract

Prokaryotic Argonaute proteins (pAgos) are homologs of eukaryotic Argonautes (eAgos) that were similarly proposed to play a role in cell defense against invaders. However, pAgos are much more diverse than eAgos and very little is known about their functional activity and target specificity *in vivo*. Here, we describe five pAgo proteins from mesophilic bacteria that act as DNA-guided DNA endonucleases and analyze their ability to target chromosomal and invader DNA. *In vitro*, the analyzed proteins use small guide DNAs for precise cleavage of single-stranded DNA at a wide range of temperatures. Upon their expression in *Escherichia coli*, all five pAgos are loaded with small DNAs preferentially produced from plasmid DNA and from chromosomal regions of replication termination. One of the tested pAgos, EmaAgo from *Exiguobacterium marinum* can induce DNA interference between multicopy sequences resulting in targeted processing of homologous plasmid and chromosomal loci. EmaAgo also protects bacteria from bacteriophage infection and is preferentially loaded with phage guide DNAs suggesting that the ability of pAgos to target multicopy elements may be crucial for their protective function. The wide spectrum of pAgo activities suggests that they may have diverse functions *in vivo* and paves the way for their use in biotechnology.

## Introduction

Prokaryotic Argonautes (pAgos) are an expanding family of programmable nucleases that use small guide oligonucleotides to recognize target nucleic acids. Initially characterized as homologs of eukaryotic Ago proteins (eAgos), which play the central role in RNA interference, they were later shown to have their own functions in cell defense against mobile genetic elements and possibly in other cellular processes (1–4).

Unlike eukaryotic Argonautes that act on RNA, most previously characterized pAgos preferentially recognize DNA targets (5–13). Most of pAgos use small DNA guides but some bind RNA guides, suggesting the existence of different routes of guide biogenesis (14,15). Interestingly, no pAgo proteins with strict specificity for RNA guides and RNA targets, characteristic to eAgos, are known. However, a new group of pAgos that use DNA guides to recognize RNA targets have been described recently, suggesting that analysis of various branches of the pAgo phylogenetic tree can lead to further discoveries of pAgos with different functional activities *in vitro* and *in vivo* (16,17).

Phylogenetic analysis revealed that pAgo proteins form three clades, long-A, long-B and short pAgos (Fig. 1A) (2,3,18). Long Agos contain six domains, N-terminal, L1, PAZ, L2, MID and PIWI (Fig. 1B), while short pAgos have the MID and PIWI domains but lack the N-terminal half. All pAgos that are predicted to be catalytically active based on the presence of a conserved catalytic tetrad in their PIWI domain belong to the long-A clade, including all experimentally characterized pAgo nucleases (Fig. 1A). In contrast, all members of the long-B and short clades contain substitutions of the key catalytic residues in the PIWI domain. These pAgos are usually co-encoded with or directly fused to nucleases and other proteins with various predicted functions (1,2,18).

**Figure 1.**
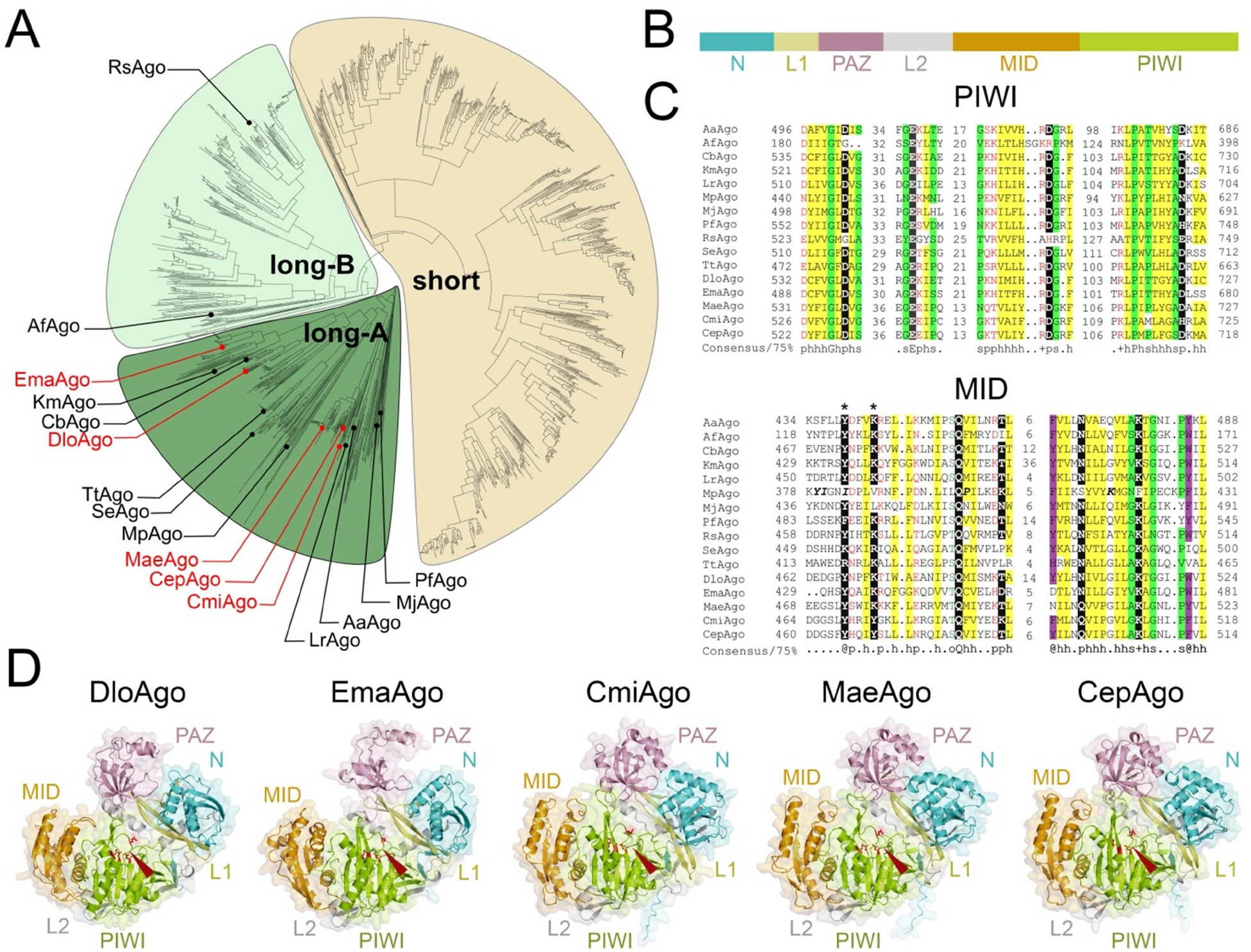
Programmable nucleases from different phylogenetic groups of pAgos. (A) Phylogenetic tree of pAgo proteins (18,19). The clades of active long pAgos (long-A), inactive long pAgos (long-B) and short pAgos are indicated. Previously studied pAgos are labeled in black. The newly characterized pAgos are shown in red. (B) Domain structure of pAgo proteins. (С) Alignment of the key elements of the catalytic site in the PIWI domain (top) and the 5’-guide-binding pocket in the MID domain (bottom) in studied pAgos. Conserved residues that play the key role in catalysis in the PIWI domain (the DEDX tetrad) or in the 5’-guide end binding in the MID domain are shown in black (the YK motif is indicated with asterisks). The 75% sequence consensus is shown underneath the alignment: ‘h’, hydrophobic residues (WFYMLIVACTH); ‘p’, polar (EDKRNQHTS); ‘s’, small (ACDGNPSTV); ‘o’, OH-containing (ST); ‘@’, aromatic (YWFH). (G) Structural models of the five studied pAgo proteins, obtained by AlphaFold. Individual domains of pAgo proteins are differently colored. The active sites are indicated with arrowheads; the catalytic tetrad residues are shown in red as stick models.

Studies of the *in vivo* functions of pAgos have been limited to just a few members of this family. We have recently shown that CbAgo from *Clostridium butyricum* can protect bacterial cells from bacteriophages (19). Several pAgos including CbAgo and TtAgo from *Thermus thermophilus* were also shown to preferentially recognize plasmid DNA during their expression in *Escherichia coli* cells, which leads to plasmid loss and decreases the plasmid transformation efficiency (8-10,19). This suggests that pAgos may have a defense function, echoed by eAgos acting in RNA interference (2,4,20). pAgos may also play alternative roles in bacteria. Thus, TtAgo from *Thermus thermophilus* was shown to participate in the completion of DNA replication by decatenating chromosomes in the presence of ciprofloxacin, and CbAgo was similarly shown to target the region of chromosomal replication termination (19,21).

Recent studies of long-B and short pAgos suggested that they can also participate in cell protection against invader genetic elements, despite the absence of intrinsic nuclease activity (22–25). Their defense function requires guide-dependent recognition of target nucleic acids and likely depends on the action of additional effector proteins, such as cellular and pAgo-associated nucleases, as well as proteins with other activities, including NADases or membrane-disrupting effectors. These effectors may lead to cell death upon pAgo-mediated target recognition and thus protect bacterial population from phages by causing abortive infection (23–26).

Here, we have analyzed predicted pAgo nucleases from several phylogenetic branches of long-A pAgos not tested in previous studies. We have shown that all of them act as DNA-guided DNA endonucleases *in vitro* and can target chromosomal and invader DNA with varying specificities *in vivo*. We have demonstrated that pAgos cooperate with the cellular double-strand break processing machinery for small DNA binding. In contrast to catalytically inactive short pAgos, their activities do not depend on the action of co-encoded effector proteins, thus enabling their action in heterologous host bacteria. One of the tested pAgos induces cleavage of multicopy elements and thus protects the cells from bacteriophage infection. The results suggest that preferential processing of multicopy elements by pAgos and cellular nucleases likely underlies the protective function of pAgos.

## Materials and Methods

The gene of EmaAgo (WP_026824436.1) was amplified by PCR from genomic DNA of *Exiguobacterium marinum* strain DSM-16307 and cloned into the pBAD-HisB plasmid. Nucleotide sequences of DloAgo (WP_055195547.1; *Dorea longicatena*), CmiAgo (WP_015159126.1; *Chamaesiphon minutus*), MaeAgo (WP_002747795.1; *Microcystis aeruginosa*) and CepAgo (WP_015201773.1; *Crinalium epipsammum*) were codon-optimized using IDT Codon Optimization Tool for expression in *E. coli*, synthesized by the IDT core facility and cloned into the pET28b expression vector (for protein purification) or into pBAD-HisB (for isolation of Ago-associated nucleic acids and analysis of phage infection) in frame with the N-terminal His_6_-tag.

For expression of EmaAgo, *E. coli* BL21(DE3) cells carrying the pBAD expression plasmid were cultivated in the LB medium with 200 µg/ml ampicillin at 37 °C until OD_600_ 0.3, cooled down to 30 °C, induced by the addition of L-arabinose to 0.1% and grown for 4 h at 30 °C. For expression of DloAgo and CmiAgo, *E. coli* BL21(DE3) was transformed with corresponding expression plasmids and the cells were cultivated in the LBN medium (Luria-Bertani medium + 0.5 M NaCl) with 0.1% glucose and 50 μg/ml kanamycin at 37 °C overnight. The overnight culture was transferred into fresh LBN supplemented with 0.1% glucose, 50 μg/ml kanamycin and 1 mM betaine, grown at 37 °C until OD_600_ 0.3-0.4, cooled down to 18 °C, induced with 0.25 mM IPTG and grown for 16 h at 18 °C. For expression of CepAgo and MaeAgo, *E. coli* BL21(DE3) cells carrying the expression plasmids were cultivated in the LB medium with 50 μg/ml kanamycin at 37 °C until OD_600_ 0.3, cooled down to 30 °C, induced with 0.1 mM IPTG and grown for 4 h at 30 °C. The cells were collected by centrifugation and stored at −80 °C.

For purification of pAgo proteins, cell pellets were resuspended in buffer A (50 mM Tris– HCl pH 7.5, 0.5 M NaCl, 20 mM imidazole, 5% glycerol) supplemented with 1 mM of PMSF and disrupted using a high-pressure homogenizer (EmulsiFlex-C5, Avestin) at 18000 psi. The lysate was cleared by centrifugation at 30,000 g for 30 min and the supernatant was loaded onto a HisTrap FF crude column (GE Healthcare) equilibrated with buffer A. The column was washed with buffer A containing 50 mM imidazole and the proteins were eluted with buffer A containing 250 mM imidazole. The eluted proteins were concentrated by Amicon Ultra Centrifugal Filter 50 kDa (Merck Millipore) and purified on a Superose 6 10/300 GL column (GE Healthcare) equilibrated with buffer GF (10 mM HEPES–NaOH pH 7.0, 0.4 M NaCl, 5% glycerol, 1 mM DTT). Fractions containing pAgo proteins were loaded onto a Heparin column (GE Healthcare) equilibrated with buffer GF, washed with 10 column volumes of the same buffer and eluted with buffer GF containing 0.7 M NaCl. The proteins were concentrated using Amicon Ultra 50 kDa, diluted in a storage buffer (10 mM HEPES-NaOH pH 7.0, 0.35 M NaCl, 50% glycerol, 0.5 mM DTT) and stored at −20 °C. The protein concentration was determined by a NanoDrop Spectrophotometer (Thermo Fischer Scientific).

Protein structure prediction was performed with AlphaFold2 algorithm (27) using ColabFold (28) (notebook AlphaFold2-mmseqs2 with default parameters). The best model for each protein was visualized with PyMol.

### Analysis of nucleic acid cleavage by pAgos

The cleavage assays were performed using synthetic guide and target DNAs and RNAs (see Table S2 for oligonucleotide sequences). In some assays, target DNAs were labeled with 3’-Cy5. Most cleavage reactions were performed in buffer containing 20 mM Tris-acetate pH 7.0, 50 mM potassium acetate, 5 % glycerol, 10 mM MnCl_2_, BSA 100 μg/ml. Plasmid cleavage by DloAgo was performed in buffer containing 10 mM HEPES-NaOH pH 7.0, 100 mM NaCl, 5% glycerol and 5 mM MnCl_2_. To analyze the effect of various divalent cations (Fig. 3A), 0.5 mM or 5 mM MgCl_2_, CoCl_2_, CuCl_2_, ZnCl_2_ or CaCl_2_ were added instead of MnCl_2_. Most cleavage assays were performed at the 5:2:1 pAgo:guide:target molar ratio at 37 °C. All guides were 5’-phosphorylated using T4 PNK (New England Biolabs) except for experiments with 5’-OH guides (Fig. 3C). For analysis of the nucleic acid specificity of pAgos (Fig. 2B), 500 nM of pAgo was mixed with 200 nM guide DNA or RNA oligonucleotides, incubated for 15 min at 37 °C for guide loading, then target DNA or RNA was added to the final concentration of 100 nM, and the reaction was stopped after 2 h incubation at 37 °C. To analyze the effects of guide length on target cleavage (Fig. 3B), the reactions were performed with 10-22 nt guides (Table S2) for 30 min for DloAgo and CmiAgo, 3 hours for CepAgo, MaeAgo and EmaAgo. To define the preference for the 5’-guide nucleotide (Fig. 3C), the reaction was carried out at 37 °C for 30 minutes for CmiAgo, 1 hour for CepAgo, EmaAgo, DloAgo, and 3 hours for MaeAgo. For analysis of temperature dependence of DNA cleavage (Fig. S2A), pAgos were loaded with G-guide DNA for 15 min at 37 °C, the samples were transferred to indicated temperatures in a water bath, corresponding target DNA was added and the samples were incubated for 15 min in the case of DloAgo and CmiAgo, for 30 min in the case of CepAgo, EmaAgo and MaeAgo. For experiments on single-turnover cleavage (Fig. S2B), 1 μM pAgo was mixed with 250 nM guide DNA and 50 nM target DNA and incubated at 37 °C for indicated time intervals. To analyze the effect of SSB proteins on ssDNA cleavage (Fig. S3), the reactions were performed at the 5:2:1:4 pAgo:guide:target:SSB molar ratio. *E. coli* SSB was purified as described before (6). Target DNA was premixed with SSB (400 nM final concentration) and incubated for 10 minutes at 37 °C. Preformed complexes of pAgos with guide DNAs was added and the reaction was carried out at 37 °C for 1, 5, 10, 15, 30 minutes. For analysis of guide-target mismatches (Fig. S4), pAgos were loaded with guide DNAs containing single-nucleotide substitutions at each position (Table S2) and incubated with the same target DNA for 1 hour (2 hours for EmaAgo) at 37 °C. All reactions were stopped after indicated time intervals by mixing the samples with equal volumes of a stop-solution (8 M urea, 20 mM EDTA, 0.005% Bromophenol Blue, 0.005% Xylene Cyanol) and treated with Proteinase K for 30 minutes at 37 °C. The cleavage products were resolved by 19% denaturing PAGE. The gels were stained with SYBR Gold (Invitrogen) or directly visualized with a Typhoon FLA 9500 scanner (GE Healthcare) (in the case of 3’-Cy5-labeled target DNA). The data were analyzed using ImageQuant (GE Healthcare), Prism 8 (GraphPad) and custom R scripts (v. 3.6.3). The data on single-turnover target cleavage were fitted to the equation Y = C + A_max_×(1-exp(-*k*_obs_ × *t*)), where Y is the efficiency of cleavage at a given time point, A_max_ is the maximum cleavage, C is the background level of cleavage, and *k*_obs_ is the observed rate constant.

**Figure 2.**
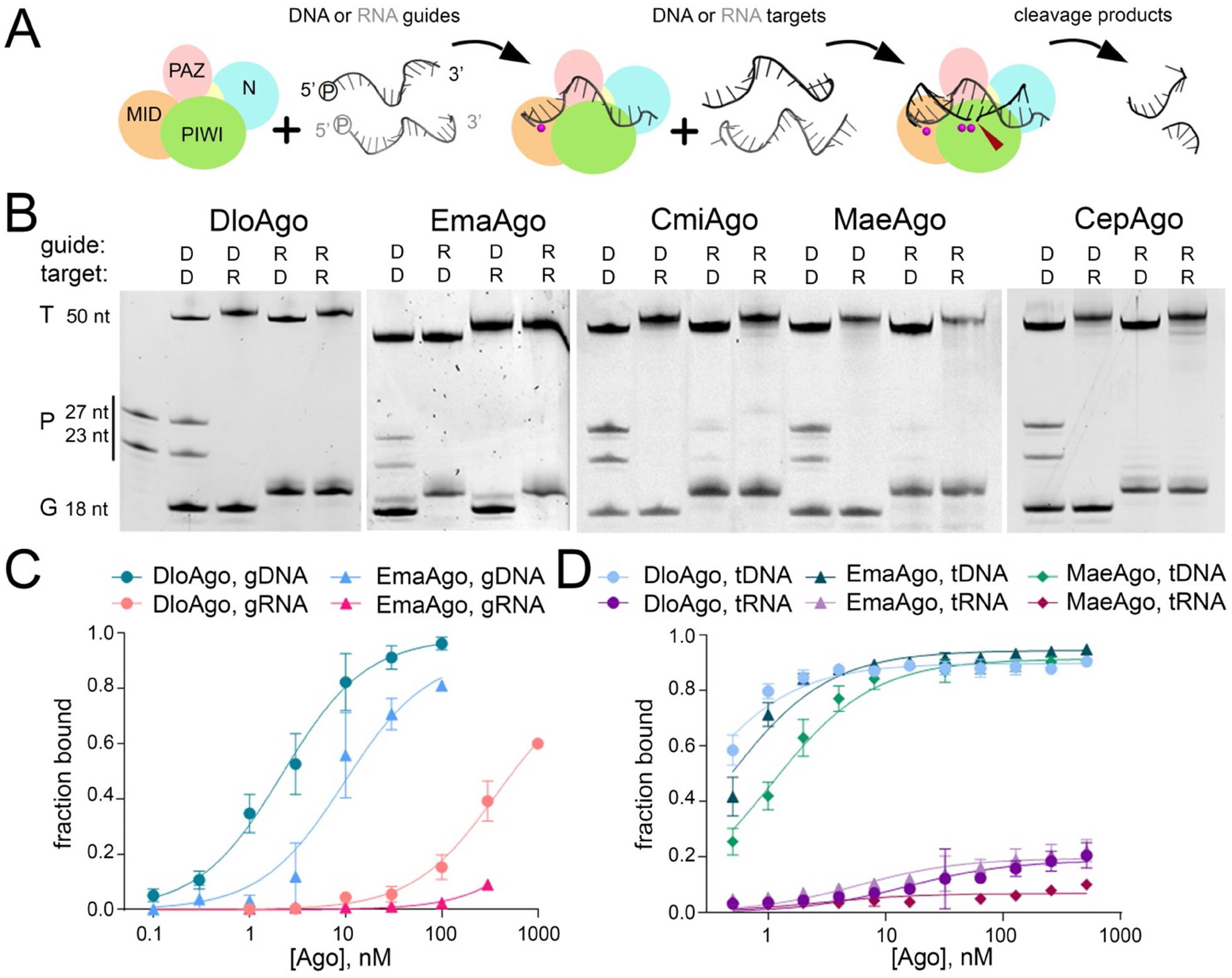
Guide and target specificity of pAgo proteins *in vitro*. (A) Scheme of the experiment. (B) Activities of studied pAgo proteins with DNA (D) or RNA (R) guides and targets. See Table S2 for oligonucleotide sequences. Positions of the target (T), guide (G) and product (P) oligonucleotides are indicated. The first lane contains the length markers (27 and 23 nt) corresponding to target cleavage between the 10^th^ and 11^th^ guide nucleotides. (C) Binding of P^32^-labeled DNA or RNA guides (gDNA, gRNA) to DloAgo and EmaAgo. (D) Binding of P^32^-labeled DNA or RNA targets (tDNA, tRNA) to DloAgo, EmaAgo and MaeAgo preloaded with guide DNA. The fraction of bound guides and targets relative to their total amount in the samples is indicated. Means and standard deviations from three independent experiments are shown.

**Figure 3.**
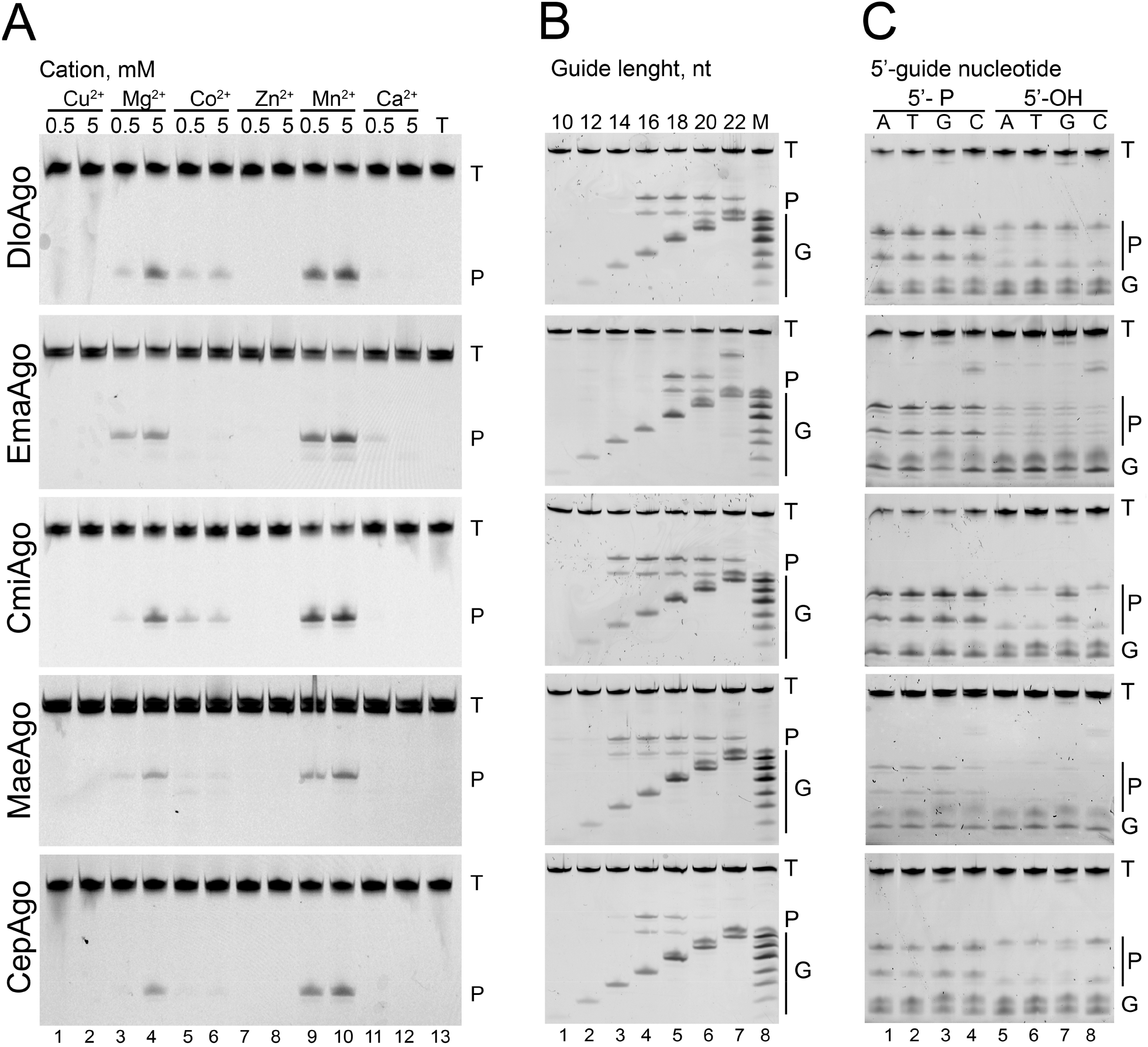
Catalytic preferences of pAgos. (A) Analysis of divalent cation dependence of DNA cleavage by pAgos. The experiments were performed with 3’-Cy5-labeled target DNA and the gels were analyzed by a fluorescence scanner. (B) Target DNA cleavage by the five pAgos proteins using guide DNAs of different lengths. The maker lane (M) contains 12, 14, 16, 18, 20, 22 and 23 nt DNA oligonucleotides. The experiments were performed with unlabeled target DNA and the gels were stained with SYBR-Gold. The experiments in panels A and B were performed with 5’-P guide DNAs. (C) Comparison of DNA cleavage with 5’-P or 5’-OH guide DNAs with various 5’-nucleotides (indicated above the gels). Positions of the target (T), guide (G) and product (P) oligonucleotides are indicated. See Materials and Methods for the reaction conditions.

For analysis of plasmid DNA cleavage, pAgo samples were separately loaded with two or four guides at 37 °C for 15 min and mixed together to the final concentration of 500 nM (see Table S2 for oligonucleotide sequences). The target plasmid pSRKKm was added to the reaction mixtures to the final concentration of 2 nM, followed by incubation for indicated time intervals at 45 °C for DloAgo and 42 °C for EmaAgo. Linear plasmids were obtained by treatment with a single cut restriction endonuclease (XhoI or KpnI, Thermo Fisher Scientific). The reactions were stopped by treatment with Proteinase K for 30 minutes at 37 °C, the samples were mixed with 6^×^ SDS-free Purple Loading Dye (New England Biolabs) supplemented with SYBR Gold, the cleavage products were resolved by native 1 % agarose gel electrophoresis and visualized with a Typhoon FLA 9500 scanner.

### Analysis of guide and target binding by pAgos

Interactions of guides and targets with pAgos were analyzed by a dot blot assay as described previously (29). For determination of apparent dissociation constants (*K*_d_) for guide binding, 5’-P^32^-labeled DNA or RNA guide oligonucleotides (0.1 nM final concentration) were mixed with increasing concentrations of pAgo in the binding buffer containing 10 mM Tris-HCl pH 7.9, 100 mM NaCl, 5 mM MgCl_2_, 5% glycerol and 100 μg/ml BSA. The mixtures were incubated for 20 min at 37°C and loaded onto a Bio-Dot Microfiltration Apparatus (Bio-Rad) loaded with nitrocellulose (0.2 μm Amersham Protran, GE Healthcare) and nylon membrane filters (Hybond N+, GE Healthcare). For analysis of target binding, 5’-P^32^-labeled target oligonucleotides (0.25 nM) were titrated with increasing concentrations of stoichiometric binary guide-pAgo complexes in the binding buffer containing 20 mM Tris-acetate pH 7.0, 100 mM potassium acetate, 5 % glycerol, 10 mM MnCl_2_, BSA 100 μg/m; the samples were incubated for 5 min at 37 °C and processed as described above. The membranes were visualized by phosphorimaging using a Typhoon FLA 9500 scanner (GE Healthcare) and analyzed by ImageQuant (GE Healthcare) and custom R scripts (v. 3.6.3). The fraction of pAgo-bound guide or target oligonucleotides was calculated as the ratio of the amounts of labeled oligonucleotides bound to the nitrocellulose membrane to the sum of the signals from the nitrocellulose and nylon membranes. The data were fitted to the equation: B = B_max_×C/(*K*_d_ + C), where B is the fraction of bound oligonucleotide, B_max_ is the maximum binding, and C is the concentration of pAgo or of the binary guide-pAgo complex.

### Analysis of the effects of pAgos on phage infection

Analysis of P1 phage infection in *E. coli* strains expressing different pAgo proteins was performed as described earlier (19). A lysate of P1vir phage from our laboratory collection was prepared from the *E. coli* MG1655 strain as previously described (19). The phage titer was determined by standard methods by counting plaque forming units (PFU). To test the effects of pAgos on cell growth during phage infection (Fig. 7A), *E. coli* strain MG1655 Z1 was transformed with pBAD plasmids encoding each of the five pAgos or a control plasmid without pAgo, grown overnight in LB with ampicillin (100 μg/ml), diluted twice with 50% glycerol, aliquoted, frozen in liquid nitrogen and stored at −80 °C. Overnight bacterial cultures were obtained from frozen aliquoted cultures and inoculated into 1 ml of fresh LB medium supplemented with CaCl_2_ (5 mM), MgSO_4_ (10 mM), ampicillin (100 μg/ml) and arabinose (0.05%) in 12-well plates. Expression of pAgos was verified by Western blot with antibodies against His-tag (Sigma) after cell growth for 2 hours at 30 °C. The phage lysate was added at various multiplicity of infection (MOI) (0.004, 0.0008, 0.00016), with a no-phage control. The MOI values were defined as the ratio of phage PFU to *E. coli* colony forming units (CFU), measured for the cell culture at the time of infection. The plates were incubated at 300 rpm at 30 °C in a CLARIOSTAR microplate reader and cell density was monitored by measuring OD_600_ every 10 min. Three independent biological replicates were performed for each strain and MOI, and means and standard deviations were calculated and plotted using a custom R script.

To determine phage titers during infection (Fig. 7B,C), 100 μl aliquots of bacterial cultures were taken at 2 and 7 hours after the start of infection and treated with 100 μl of chloroform for 15 s with vortexing. The aqueous phase was transferred to a fresh tube and stored at 4 °C for no more than 5 days. Serial dilutions were prepared in 10 mM MgSO_4_, 5 mM CaCl_2_ and aliquots were plated on LB plates covered with top agar containing 10 mM MgSO_4_, 5 mM CaCl_2_ and MG1655 Z1 cells grown until OD_600_ of 0.2. Phage plaques were counted after overnight growth at 30 °C.

To measure the content of phage DNA at different stages of infection (Fig. 8), bacterial cultures were infected with phage P1vir at MOI=7×10^−5^ as described above in 24 well plates and were grown at 200 rpm at 30 °C in a CLARIOSTAR microplate reader. Aliquots of cell culture (100 μl) were taken every 30 minutes. The cells were collected centrifugation, and bacterial pellets were washed twice by resuspending in 0.3 ml of cold saline solution (0.9% NaCl) followed by centrifugation. The cells were lysed by heating at 98 °C for 10 minutes in 50 μl of milliQ water. Cell debris was precipitated by centrifugation in an Eppendorf Centrifuge 5424 at 14,680 rpm for 10 min, supernatant was transferred to new tubes and several tenfold dilutions were used for quantitative PCR. To estimate the relative abundance of bacterial and phage DNA, specific primers were designed for bacterial and phage genomes using Primer-BLAST (30). The absence of primer dimers was verified by ThermoFisher Multiple Primer Analyser. The melting curve and efficiency plots were used to validate the specificity and efficiency of each primer pair. Quantitative PCR was performed using qPCRmix-HS with SYBR Green 1 premix (Evrogene) in a Bio-Rad Real-Time CFX96 Touch thermal cycler using the following conditions: initial denaturation at 95 °C for 3 minutes, 40 cycles of denaturation at 95 °C for 10 seconds, annealing at 62 °C for 15 seconds and synthesis at 72 °C for 15 seconds. The ratio of phage and cell DNA at different time points was calculated and plotted using a custom Python script.

### Preparation and analysis of small DNA and genomic DNA libraries

For isolation of pAgo-associated nucleic acids, *E. coli* BL21(DE3) strains carrying the expression plasmids with pAgos (pBAD-HisB) were cultivated in the LB media with 200 μg/ml ampicillin at 37 °C until OD_600_ 0.3, cooled down to 30 °C, induced by the addition of L-arabinose to 0.1% and grown for 4 h at 30 °C. The cells were collected by centrifugation and disrupted with a high-pressure homogenizer. pAgos were pulled down using Co^2+^-Talon Metal Affinity Resin (Takara) as described previously (19). Eluted proteins were treated with Proteinase K for 30 minutes at 37 °C, small nucleic acids were extracted with phenol-chloroform, ethanol-precipitated, dissolved in water, treated with DNase I or RNase A and analyzed by PAGE as described (19).

Libraries for high-throughput sequencing of small DNAs were prepared according to the previously published splinted ligation protocol (19). Briefly, nucleic acids extracted from pAgos were treated with RNase A (Thermo Fisher), purified by PAGE, small DNAs (14-20 nt) were eluted from the gel in 0.4 M NaCl overnight at 21 °C, ethanol precipitated, dissolved in water, phosphorylated with polynucleotide kinase (New England Biolabs), and ligated with adaptor oligonucleotides using bridge oligonucleotides as described in (19). The ligated DNA fragments were purified by denaturing PAGE, amplified and indexed by the standard protocol for small RNA sequencing (New England Biolabs). Small DNA libraries were sequenced using the HiSeq2500 platform (Illumina) in the rapid run mode (50-nucleotide single-end reads).

For DNA sequencing during bacteriophage infection, *E. coli* strain MG1655 Z1 was transformed with an empty pBAD plasmid or pBAD encoding EmaAgo. Overnight cell cultures (2.5 ml, 8.3×10^9^ CFU/ml) were inoculated into 0.5 liter of LB supplemented with 10 mM MgSO_4_, 5 mM CaCl_2_, 0.05% L-arabinose and 100 μg/ml ampicillin. The P1vir lysate (100 μl, 3×10^8^ PFU/ml, MOI=0.0015) was added and the cultures were grown for 2.5 hours at 200 rpm at 30 °C. From each culture, 10 ml aliquots were taken, centrifuged, and the cell pellets were washed twice with 0.9% ice-cold NaCl, aliquoted and frozen. The cells from the rest volume were also pelleted, resuspended in ice-cold buffer containing 30 mM Tris-HCl, 200 mM NaCl, 5% glycerol, 2 mM PMSF and lysed using a high-pressure cell homogenizer. Lysates were clarified by centrifugation and 200 μl of Co^2+^ resin (TALON) equilibrated with the same buffer was added. The samples were incubated at 4 °C for 2 hours with rotation. The affinity resin was precipitated, washed 3 times with 800 μl of the same ice-cold buffer without PMSF and then with the same buffer with 5 mM imidazole. Argonaute proteins were eluted with 500 μl of the same buffer supplemented with 300 mM imidazole. The extracted proteins were used for small DNA purification by phenol-chloroform extraction and ethanol precipitation.

For genomic DNA sequencing, total DNA from infected *E. coli* cultures was extracted from the aliquots taken from the same *E. coli* cultures (2.5 hours after infection at MOI=0.0015). Genomic DNA libraries were prepared using the NEBNext Ultra II FS DNA Library Prep Kit (NEB), with the insert size in the range of 150 to 250 bp. Barcodes were introduced to both DNA ends during library amplification with NEBNext multiplex oligos for Illumina (NEB). Genomic DNA libraries were sequenced using HiSeq2500 (Illumina) with 200 nt single-end reads. The complete genomic sequence of the P1 phage used in our experiments was assembled from contigs obtained from the same library and the remaining gaps were closed by Sanger sequencing. Small DNA libraries were prepared from the same samples of infected cells in the same way as described above for non-infected cultures. The list of all analyzed genomic and small DNA libraries is shown in Supplementary Table S3.

Analysis of small DNA sequences was performed as described previously (19). Briefly, raw reads were quality checked with FastQC (v. 0.11.9), adaptors were trimmed and the reads with length less than 14 nt were discarded from further analysis with CutAdapt (v. 2.8). Reads were aligned to the reference genomic DNA (Refseq accession number NC_012971.2 for BL21(DE3), NC_000913.3 for MG1655, GeneBank number OP279344 for phage P1) allowing no mismatches, using Bowtie (v. 1.2.3). Calculation of genome coverage was made using BEDTools (v. 2.27.1) and custom Python scripts. Plasmid coverage was calculated as a rolling mean (200 nt window, 10 nt step). Genomic and phage DNA coverages were calculated after removing reads that aligned to more than one DNA molecule and plotted in 1000 nt windows. The ratio between small DNAs mapped to the plus-strand and minus-strand was calculated as a rolling mean (50 kb window, 10 kb step). To build metaplots of small DNA distribution around Chi sites in the genome, the region of replication termination (1.2-1.7 Mb chromosomal coordinates) was discarded from the analysis. 20 kb regions centered around Chi-sites were split into 500 nt bins. Bins coverage was calculated separately for each strand and averaged. Nucleotide Logos were generated using custom python scripts and ggseqlogo package for R (v. 0.1) (31). Only reads with the length >16 nt were taken into analysis. Reads longer than 17 nt were truncated from the 3’-end to 17 nt. All plots were generated in R (v. 3.6.3) using custom scripts.

## Results

### Identification of candidate pAgo nucleases from mesophilic bacteria

To explore new pAgo nucleases, we selected several candidate proteins from different branches of the long-A clade, including DloAgo from *Dorea longicatena*, EmaAgo from *Exiguobacterium marinum*, MaeAgo from *Microcystis aeruginosa*, CmiAgo from *Chamaesiphon minutus* and CepAgo from *Crinalium epipsammum* (Fig. 1A). All these pAgos are encoded by mesophilic nonpathogenic bacteria and it could therefore be expected that they should be active at ambient temperatures.

Alignment of the newly selected pAgo proteins indicates that all of them have a canonical 6 domain structure (Fig. 1B) and contain the catalytic tetrad residues in the PIWI domain, DEDD in DloAgo, EmaAgo, MaeAgo and CepAgo and DEDH in CmiAgo (Fig. 1C). They also contain a conserved binding pocket for the 5’-end of a guide molecule in their MID domains, including the YK motif (or YY in CmiAgo and CepAgo) directly involved in the 5’-nucleotide interactions (Fig. 1C). Furthermore, structural modeling suggests that the overall architecture of the candidate pAgos is similar to previously studied DNA-guided DNA nucleases, such as TtAgo, CbAgo and MjAgo (Fig. 1D). At the same time, these proteins contain substitutions of non-catalytic residues in the active site and the MID pocket and also variations in other protein parts, which may potentially affect their activities (Fig. 1C). To get insight into the functional differences of these pAgos, we characterized their activities *in vitro*, using purified proteins, and *in vivo*, during their expression in the heterologous *E. coli* system.

### Novel pAgo proteins act as DNA-guided DNA nucleases

The genes of the five pAgo proteins were obtained by chemical synthesis after codon-optimization for expression in *E. coli* or obtained by PCR from genomic DNA of host bacteria, cloned into expression vectors and expressed in *E. coli* (see Materials and Methods). The proteins were purified by Ni^2+^-affinity, gel-filtration and heparin-affinity chromatography (Fig. S1).

To characterize the nucleic acid specificity of pAgos, we loaded them with DNA or RNA guides and analyzed cleavage of single-stranded DNA and RNA targets containing regions of complementarity to the guide molecules (Fig. 2A). We found that all five pAgos have DNA-guided DNA endonuclease activity and cut their DNA targets at the expected position between the 10^th^ and 11^th^ guide nucleotides (Fig. 2B). Low level of DNA cleavage is also observed with RNA guides for MaeAgo and CmiAgo (Fig. 2B), while none of the pAgos can cut RNA with either DNA or RNA guides under these conditions.

To explain the observed specificity, we analyzed the interactions of pAgos with guides and targets using a dot-blot assay. Comparison of the affinities of DloAgo and EmaAgo for DNA and RNA guides of the same sequence showed that DNA is bound much stronger than RNA (apparent *K*_d_ values of 2.4±0.6 nM and 11.4±6 nM, respectively, for DNA guides and >100 nM for RNA guides; Fig. 2C), in agreement with the observed specificity of the cleavage reaction for DNA guides. We further compared interactions of pAgos loaded with DNA guides with DNA and RNA targets. It was found that three tested pAgos all have very high affinity for DNA targets (apparent *K*_d_s of 0.25±0.05, 0.48±0.06 and 1.3±0.2 nM for DloAgo, EmaAgo and MaeAgo, respectively) (Fig. 2D). In contrast, they do not form complexes with RNA targets (*K*_d_s>500 nM). Therefore, the absence of RNA cleavage by these pAgos is likely explained by their preference for DNA targets and inability to stably bind RNA targets.

### Requirements for specific DNA cleavage by pAgos *in vitro*

Divalent cations are essential for nucleic acid cleavage by Ago proteins. Most previously studied proteins can use either Mg^2+^ or Mn^2+^ as cofactors. We tested the ability of our set of pAgos to cleave DNA in the presence of Mg^2+^, Mn^2+^, Co^2+^, Cu^2+^, Zn^2+^ and Ca^2+^. For all five proteins, the highest activity was observed with Mn^2+^, and all were also active with Mg^2+^ (Fig. 3A). For some pAgos (DloAgo and CmiAgo), weak DNA cleavage was also observed in the presence of Co^2+^. No activity was detected in the presence of Cu^2+^, Zn^2+^ or Ca^2+^ for any pAgo. Thus, further *in vitro* experiments were performed in the presence of Mn^2+^.

The ability of pAgos to interact with diverse guide molecules may be essential for their *in vivo* functions and for their potential use as programmable nucleases. Prokaryotic and eukaryotic Ago proteins are known to use guide nucleic acids of varying length, and some of them have specific preferences for the 5’-guide nucleotides. Our set of pAgos can use DNA guides ranging from 14 to 22 nucleotides for various proteins (Fig. 3B). Substitutions of the 5’-guide nucleotide do not affect the efficiency of cleavage, suggesting that this nucleotide is not specifically recognized by these pAgos (Fig. 3C, lanes 1-4). Most previously described Ago proteins preferentially use 5’-phosphorylated guide nucleic acids. The presence of the 5’-phosphate in guide DNA is also important for precise and efficient target cleavage by the five new pAgos. When the reactions are performed with non-phosphorylated 5’-OH guide DNAs, the efficiency of cleavage is decreased for all pAgos (Fig. 3C, lanes 5-8). In most reactions with 5’-OH guide DNAs, the position of cleavage is also shifted by one nucleotide toward the guide 3’-end, so that the target is cut between the 11^th^ and 12^th^ guide positions (Fig. 3C, lanes 5-8).

When loaded with optimal guide DNA, pAgo proteins can cleave DNA at moderate temperatures starting from 18 °C (DloAgo, EmaAgo, CmiAgo) or 25 °C (MaeAgo, CepAgo), in agreement with the mesophilic nature of their host bacteria (Fig. S2A). Remarkably, all five proteins remain active up to 60 °C, and EmaAgo still retains activity at 65 °C. Thus, these pAgo proteins can be used as programmable nucleases in a wide range of temperatures. To compare the catalytic rates of pAgos, we measured the kinetics of single-stranded DNA cleavage under single-round conditions (when preformed pAgo-guide DNA complex was present in excess over the target) at 37 °C, the physiological temperature at which all five pAgos are active. It was found that the reaction rates are different for various pAgos, with half-times of the reaction varying from ~10 to 150 min, in the range of activities previously reported for pAgo proteins (Fig. S2B).

In our experiments, only a fraction of the single-stranded DNA target was usually cleaved during the reaction. The incomplete cleavage might be explained by the formation of nonproductive pAgo-DNA complexes or by the presence of secondary structures in the target DNA thus making it inaccessible for cleavage. To test this hypothesis, we analyzed whether the cleavage efficiency can be enhanced in the presence of a single-stranded DNA binding protein, *E. coli* SSB, which could help to unfold inhibitory secondary DNA structures. It was found that for three pAgos (EmaAgo, MaeAgo and CepAgo), SSB indeed significantly increased the efficiency of DNA cleavage (Fig. S3).

The specificity of target recognition is an important characteristic of programmable nucleases, which is especially important for their practical applications. We tested the effects of single-nucleotide mismatches between the guide and target oligonucleotides on the activity of pAgos. In contrast to eukaryotic Agos, mismatches in the seed region of guide DNA (positions 2-8) have only mild or no effects on target DNA cleavage by the analyzed pAgos (Fig. S4). Mismatches in the central guide region surrounding the cleavage site (positions 9-12) do not prevent target DNA cleavage by DloAgo, EmaAgo and CmiAgo but inhibit target cleavage by MaeAgo and CepAgo. Mismatch at position 10 also shifts the cleavage position for DloAgo to between 11^th^ and 12^th^ guide nucleotides. Mismatches in the 3’-supplementary region have the strongest effects on target cleavage by various pAgos. Mismatches at positions 14-15 decrease cleavage by EmaAgo, MaeAgo, and CepAgo while mismatch at position 13 inhibits the activity of all pAgos (Fig. S4).

Therefore, the new pAgos can site-specifically cleave DNA targets in a wide range of conditions when programmed with small 5’-phosphoryated DNA guides, and the full guide-target complementarity is important for cleavage.

### Plasmid DNA cleavage by pAgos

Previous experiments with various Argonaute proteins demonstrated that single-stranded DNA or RNA is a preferred substrate for guide-directed cleavage while double-stranded nucleic acids are cleaved less efficiently. Specific cleavage of plasmid DNA could be observed with some pAgos in AT-rich target regions and at elevated temperatures (5,9,11,32,33). Some pAgos were also shown to process double-stranded DNA in the absence of guides, resulting in its “chopping” into smaller fragments that can possibly be used by pAgos as guides during subsequent rounds of DNA cleavage (5,33). To test whether pAgos analyzed in this study can process double-stranded DNA, we measured the activities of DloAgo and EmaAgo with a supercoiled plasmid at 42-45 °C. The reaction was performed in the absence or in the presence of one or two pairs of guide DNAs corresponding to two different sites of the plasmid, so that DNA cleavage with any pair of these guides would result in plasmid linearization and cleavage with the two pairs would produce two DNA fragments of ~1 and 4.8 kilobases (Fig. 4A,B).

**Figure 4.**
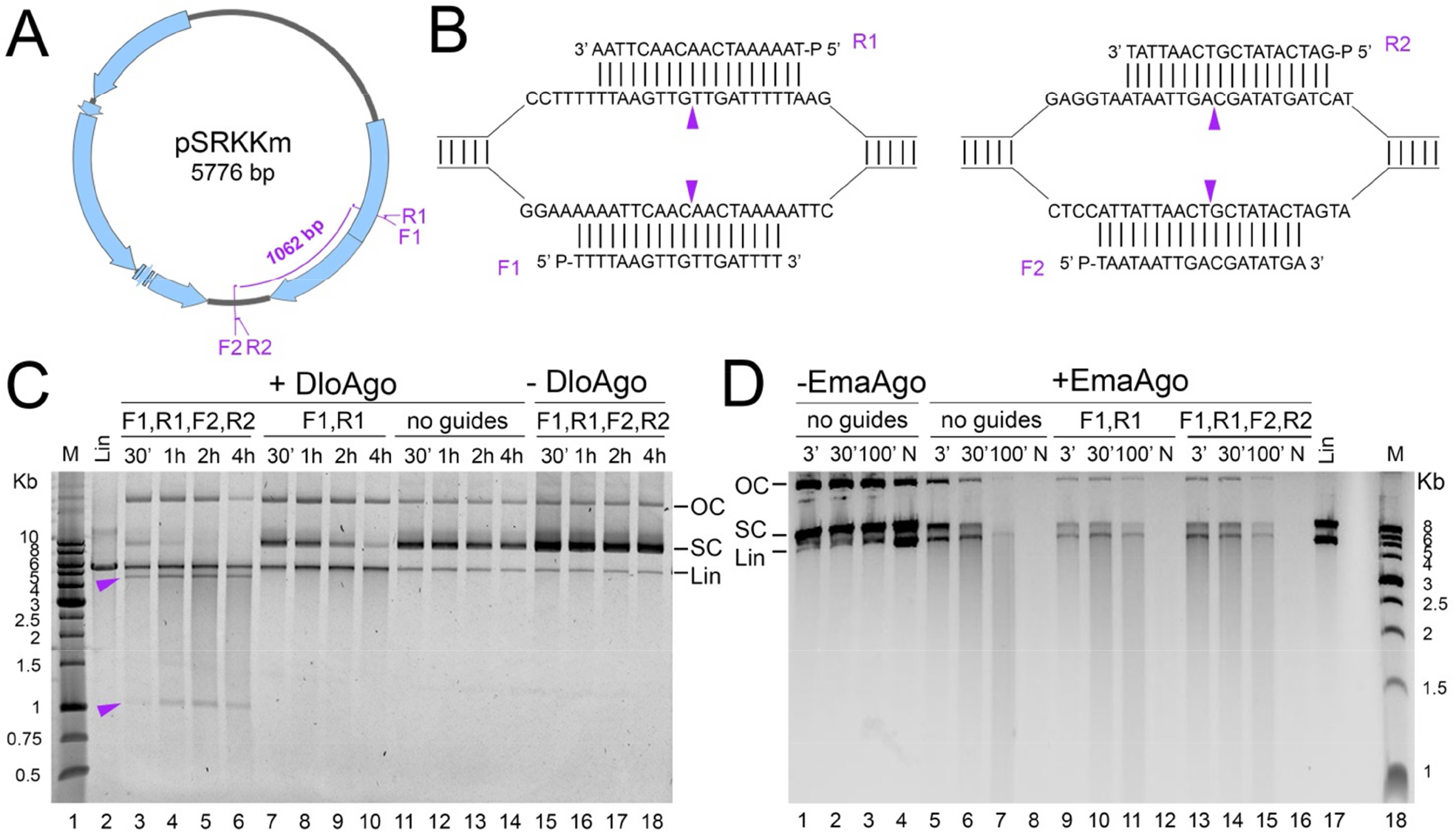
Analysis of plasmid DNA cleavage by DloAgo and EmaAgo. (A) Scheme of the target plasmid. Positions of guide DNAs (F1, R1, F2, R2) are indicated. (B) Sequences of guide DNAs and corresponding target plasmid regions. Expected cleavage sites are indicated with arrowheads. (C) Plasmid cleavage by DloAgo (at 45 °C). (D) Plasmid cleavage by EmaAgo (at 42 °C). The cleavage was performed for indicated time intervals (N, overnight for 18 hours) with empty pAgos or with pAgos independently loaded with two or four guide DNAs. Positions of supercoiled (SC), relaxed open circle (OC) and linear (Lin) plasmid DNA forms are shown. Positions of the expected cleavage products are indicated with arrowheads. In control reactions, the plasmid was linearized using KpnI (C) or XhoI (D) restriction endonucleases.

The activities of DloAgo and EmaAgo in this reaction were remarkably different. DloAgo did not process the plasmid in the absence of guide DNAs (Fig. 4C, lanes 11-14), linearized it in the presence of two guide DNAs (lanes 7-10) and cut it into two fragments of the expected lengths in the presence of four guide DNAs (lanes 3-6). However, this reaction was not complete even within 4 hours and nonspecific cleavage was additionally observed at long time intervals, resulting in DNA smearing (*e.g.* lanes 5, 6 and 9, 10). In contrast, incubation of EmaAgo with the plasmid resulted in its cleavage into the smear of nonspecific DNA products of various lengths independently of the presence of guide DNAs in the reaction, resulting in complete plasmid degradation after overnight incubation (Fig. 4D, lanes 5-8, 9-12 and 13-16).

### Preferences for nucleic acid binding by pAgos *in vivo*

Our next goal was to determine the spectrum of *in vivo* activities of the pAgo proteins. To reveal their nucleic acid specificity in bacterial cells, we analyzed nucleic acids associated with each of the five pAgos when they were expressed in *E. coli* (Fig. S5A). Nucleic acids isolated from pAgos after one-step purification by metal-affinity chromatography were treated with either DNase or RNase to determine whether the bound fraction is DNA or RNA and analyzed by gel-electrophoresis. It was found that all five pAgos are associated with small DNAs (smDNAs) that range in size from 12 to 24 nucleotides for various proteins (Fig. S5C). Therefore, all five pAgos bind small guide DNAs *in vivo*, in agreement with their specificity observed *in vitro*.

To determine the repertoire of pAgos-associated DNAs, we performed high-throughput sequencing of smDNA libraries obtained from each pAgo. The resulting reads were mapped to the *E. coli* chromosome and the plasmid used for pAgo expression. The length distribution of sequenced smDNAs corresponded to the range of smDNAs bound to pAgos (with the maximal length of 24, 19, 20, 19 and 18 nucleotides for DloAgo, EmaAgo, CmiAgo, MaeAgo and CepAgo, respectively) (Fig. S6A). No strong nucleotide biases were observed along the smDNA sequences for any pAgo (with weak preference for 5’-G for MaeAgo and CepAgo) (Fig. S6B), consistent with the absence of 5’-nucleotide specificity during DNA cleavage *in vitro* (Fig. 3C). Analysis of GC-content along the smDNA sequences also demonstrated only minor variations in nucleotide composition of guide DNAs for various pAgos. The most prominent trend was an increased GC-content in the seed region and around the cleavage site for CmiAgo and MaeAgo (Fig. S6C). At the same time, the average GC-content of guide DNAs was similar to the GC-content of genomic DNA in *E. coli* (~49-53% for four pAgos, 56.6% for CmiAgo). Overall, this analysis demonstrated that the five pAgo proteins can interact with a wide range of guide smDNAs without obvious sequence-specificity.

### Different modes of genomic DNA targeting by pAgos

To reveal the preferences of pAgos for plasmid or chromosomal DNA, we compared the numbers of smDNAs associated with each pAgo corresponding to the expression plasmid and the chromosome. Plasmid-derived smDNAs were strongly enriched in all five pAgo proteins (Fig. 5A). When the read numbers were normalized by the relative lengths of the plasmid and chromosomal DNA and the plasmid copy number (~12 for pBAD), there were 7-20 times more plasmid-derived smDNAs than expected in the case of random sampling of cellular DNA by pAgos. For all pAgos, the reads were distributed throughout the entire plasmid sequence indicating that the whole plasmid is processed into smDNAs loaded into pAgos (Fig. 5A). The patterns of plasmid DNA coverage were different for different pAgos, indicating that plasmid DNA is differently targeted by pAgos. Most efficient smDNA processing was observed for EmaAgo. Some preference for the plasmid origin of replication was observed for DloAgo, EmaAgo and CmiAgo (Fig. 5A).

**Figure 5.**
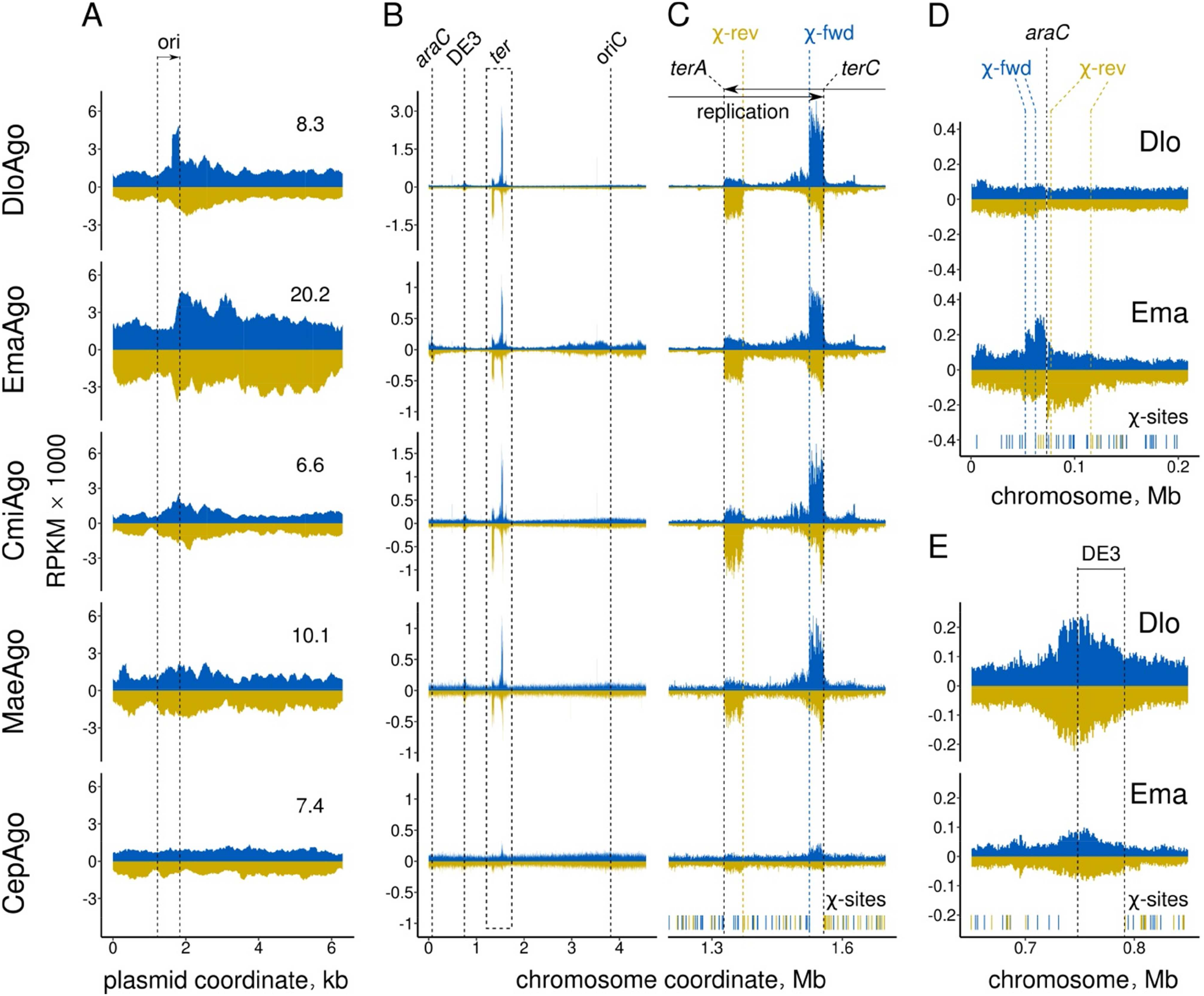
Analysis of bacterial DNA targeting by pAgos. (A) Targeting of plasmid DNA by pAgos. SmDNA coverage of the pBAD expression plasmid for the five pAgos proteins (shown in a 200-nucleotide moving window for the plus and minus DNA strands; blue and olive, respectively). The enrichment of plasmid-derived smDNAs relative to the expected coverage (based on random sampling of genomic and plasmid DNA with normalization to the plasmid copy number) is shown on each plot. (B) Distribution of smDNAs associated with pAgos along the *E. coli* BL21(DE3) chromosomal sequence shown individually for the plus (blue) and minus (olive) genomic DNA strands. Positions of *oriC*, *terA*, *terC*, *araC*, the DE3 prophage, and the directions of replichores are indicated. (C)-(E) Patterns of DNA targeting at individual chromosomal regions for various pAgos: the *ter* region (C), the *araC* gene (D) and the DE3 prophage (E). Positions of Chi sites in the plus (blue) and minus (olive) genomic strands below the plots. For each region, the closest Chi sites oriented toward the target site in each strand are shown with dotted lines (blue, Chi sites in the plus strand leftward of the target site; olive, Chi sites in the minus strand rightward of the target site). Small DNA density is shown as RPKM (reads per kilobase per million reads in the smDNA library) in 1 kilobase windows.

We then analyzed the chromosomal distribution of smDNAs associated with each pAgo protein (Fig. 5B, Fig. S7A). For all five pAgos, smDNAs were derived from both DNA strands from the entire chromosome, with several peaks of preferential smDNA processing, which included the replication termination (*ter*) region, the DE3 prophage and the *araC* gene (observed only with EmaAgo) (Fig. 5B).

To understand the details of smDNA biogenesis, we took a closer look into these genetic regions. For all pAgos, the most prominent smDNA peaks can be observed at the sites of replication termination, *terA* and *terC* (for CepAgo, these peaks are small but still detectable) (Fig. 5B,C). The size of the peak at *terC* is higher than that at *terA*, which corresponds to the shorter distance between the origin of replication and *terC* and the higher frequency of replication termination at *terC* (Fig. 5C) (34). In addition, small peaks at the next pair of the replication termination sites, *terD* and *terB*, can be detected for some pAgos, likely resulting from the replication readthrough at the innermost sites *terA* and *terC* (Fig. 5C and S7B).

Termination of replication at *ter* sites is accompanied by formation of double-stranded DNA ends, which are further recognized by the RecBCD machinery and repaired (35–37). RecBCD is a helicase-nuclease that processes double-stranded ends until the recognition of a Chi sequence (5'-GCTGGTGG-3', the 3’-end of which should be oriented toward the processed DNA end) and then loads RecA on the 3’-terminated strand, thus initiating homologous recombination (38–40). For all pAgos, the peaks of smDNAs in the termination region are confined between the *ter* sites and the closest Chi sites oriented toward the corresponding *ter* site (Fig. 5C). This indicates that these smDNAs are likely produced by RecBCD after replisome stalling at *ter* sites, until it stops DNA processing after the recognition of Chi.

At the *ter* sites, there is a prominent strand asymmetry of smDNA loading for all five pAgos, with the peaks at *terC* and *terA* enriched with smDNAs derived from the plus and minus genomic strands, respectively, which correspond to the 3’-terminated strands at these *ter* sites (Fig. 5C). This further suggests that smDNAs are produced in an asymmetric way during processing of double-stranded DNA ends by RecBCD.

Another peak of smDNAs detected for all pAgo proteins was located at the region of the lambda DE3 prophage present in the *E. coli* strain used for pAgo expression (Fig. 5E). The center of the peak corresponded exactly to the left end of the prophage suggesting that chromosomal DNA processing in this region might be caused by prophage excision (see Discussion).

EmaAgo, but not other pAgo proteins also targeted the chromosomal region around the *araC* gene (Fig. 5B,D). This peak of smDNAs likely results from DNA interference between the plasmid and chromosomal copies of *araC* (which is the only chromosomal gene present in the plasmid used for expression of pAgos). In addition, smDNAs bound to EmaAgo but not to other pAgos were enriched around rRNA operons and IS elements, clustered around the origin of replication (Fig. S7B). These additional peaks likely result from DNA interference between multicopy elements present in the chromosome.

The chromosomal peak around *araC* includes about 20 upstream and 40 kb downstream from *araC* indicating extensive smDNA processing in this region (Fig. 5D). Similarly to the *ter* sites, the distribution of smDNAs between the two genomic strands around *araC* is asymmetric. From both sides, larger amounts of smDNAs are produced from one of the strands, the 3’-end of which is oriented toward *araC* (“top” strand from the left side of *araC* and “bottom” strand from the right side of it). Furthermore, the amounts of smDNAs are strongly decreased after the first Chi site oriented toward *araC* in each strand (Fig. 5D). This indicates that EmaAgo loaded with plasmid-derived guides can induce double-strand breaks in the chromosomal *araC* locus, which are further processed by RecBCD generating smDNAs bound by EmaAgo.

To reveal whether the generation of smDNAs bound to pAgos depends on RecBCD we analyzed the distribution of smDNAs around Chi sites on the whole chromosome (excluding the *ter* region between *terA* and *terC*). Analysis of metaplots built from averaging of multiple Chi sites showed pronounced differences between the proteins. For EmaAgo, the amounts of smDNAs produced from the DNA strand co-oriented with the Chi sequence (5’-GCTGGTGG-3’) are highly asymmetric and drop from the 5’-end of Chi (orange, Fig. 5F). This pattern can be explained by participation of RecBCD in the smDNA processing, starting at double-strand breaks along the chromosome (which may be spontaneously formed during replication) and going in the upstream direction (3’ to 5’) along the 3’-terminated strand until Chi. In contrast, no strong asymmetry in smDNAs distribution is found for the opposite DNA strand, the 5’-terminus of which is oriented toward the double-strand break (gray, Fig. 5F). A similar but less pronounced asymmetry in the processing of two DNA strands is observed for DloAgo (Fig. 5F). At the same time, no significant differences in the amounts of smDNAs at the two sides of Chi sites are seen for the remaining three proteins, CmiAgo, MaeAgo and CepAgo (Fig. 5F). This indicates that RecBCD may not participate in smDNA processing for these proteins (except for the *ter* region, Fig. 5C), and its role may be taken by other nucleases.

Finally, to reveal possible preferences of the pAgo proteins for a particular DNA strand during DNA replication, we calculated the ratio of smDNAs generated from the plus and minus genomic strands along the chromosome (Fig. 5G). As expected, strong preferences can be observed for the minus and plus strands at *terA* and *terC*, respectively, for all pAgos, resulting from asymmetric DNA processing at *ter* sites (see above). However, strand-specific smDNA distribution in other chromosomal regions is different for various pAgos. For DloAgo, the leading strand (the 3’-end of which is co-oriented with the direction of replication) is preferentially targeted for both replichores (the ratio is >1 for the rightward replisome and <1 for the leftward replisome) (Fig. 5G). In contrast, for EmaAgo and CepAgo, there is a significant bias for smDNAs corresponding to the lagging DNA strand (the 3’-end of which is oriented in the opposite direction relative to replication) for both replichores (Fig. 5G). Finally, for CmiAgo and MaeAgo, this ratio is close to 1 along the whole chromosome except the *ter* sites (Fig. 5G). Therefore, different pAgo proteins may have different preferences for replication intermediates during smDNA biogenesis.

### Effects of pAgos on bacteriophage infection

To explore possible functions of the pAgo proteins in cell defense against phages, we tested whether they can protect *E. coli* cells from infection with bacteriophage P1vir, a lytic variant of phage P1 with a double-stranded DNA genome. *E. coli* strains expressing each of the five pAgo proteins or containing an empty expression plasmid were infected with phage P1 at various multiplicities of infection (MOI) in liquid medium, and cell density was monitored over time. Western blot analysis demonstrated that all five proteins were expressed at comparable levels in these conditions (Fig. S8). In the absence of phage, all strains grown with the same rate (Fig. 6A, left). Even at the lowest tested MOI (0.00016), the growth of the control strain was halted at ~6 hours post infection, as a result of phage-induced cell lysis. The growth of strains expressing DloAgo, CmiAgo, MaeAgo or CepAgo was similarly inhibited. In contrast, expression of EmaAgo protected the cells from infection under these conditions (Fig. 6A). At medium MOI (0.0008), the growth of all strains except the strain expressing EmaAgo was severely affected, and the optical densities of all the cultures dropped to the starting level after ~6 hours post infection. In a sharp contrast, the strain expressing EmaAgo continued growth under these conditions. At the highest MOI (0.004), the growth of all strains was severely affected but EmaAgo still provided some level of protection (Fig. 6A).

**Figure 6.**
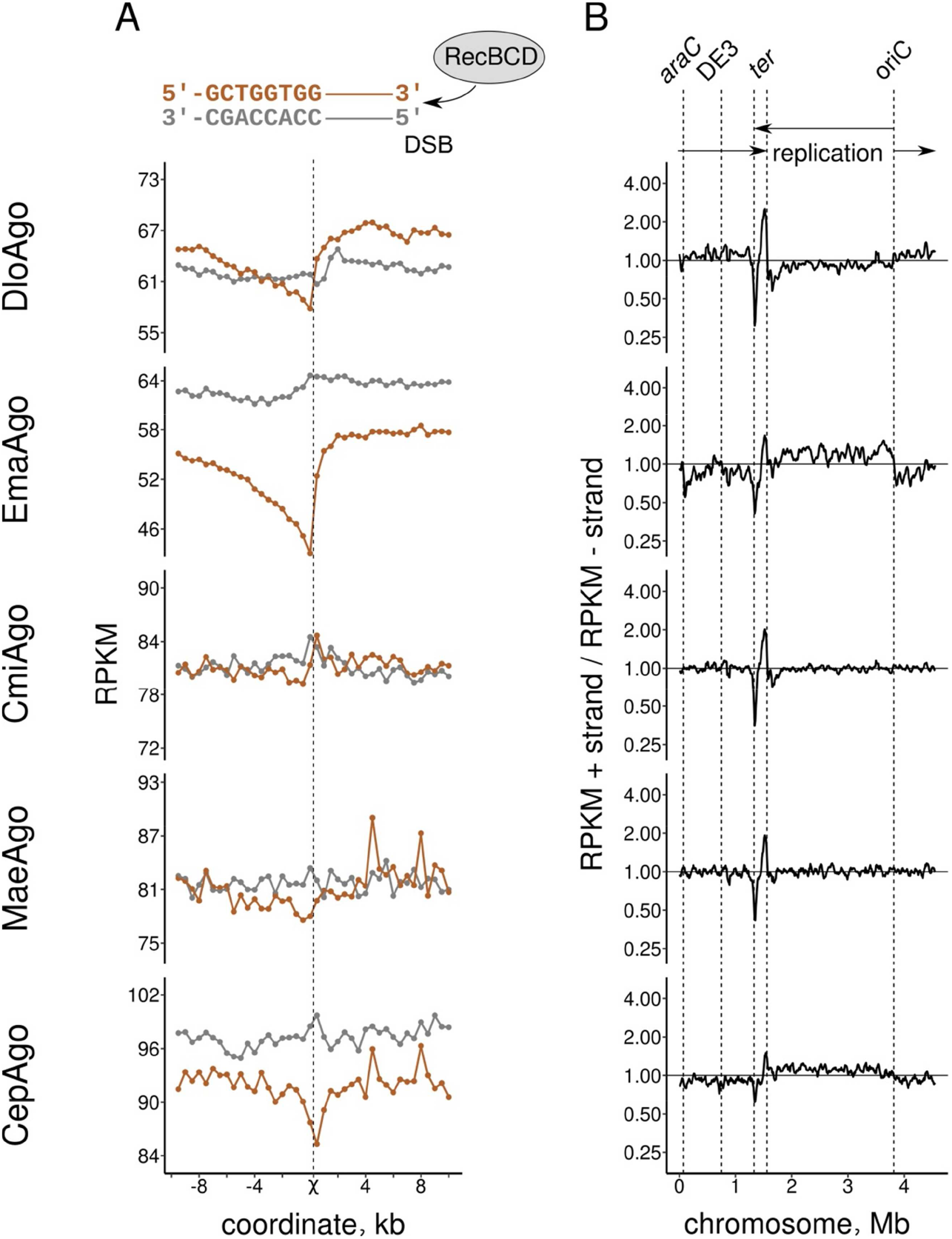
Whole-genome biases in smDNAs associated with pAgos. (F) Metaplot of the densities of smDNAs generated from the two DNA strands around Chi sites. Orange lines show smDNAs from the DNA strand co-oriented with the Chi sequence in the 5’-3’ direction (5’-GCTGGTGG-3’). Gray lines show smDNAs from the oppositely oriented DNA strand. The data were averaged for all Chi sites in both genomic strands except for the *ter* region (1.2-1.7 Mb chromosomal coordinates; 429 out of 482 sites in the plus strand and 460 out of 496 sites in the minus strand) in 0.5 kb windows. (G) Asymmetry in the chromosomal distribution of smDNAs for various pAgos. The ratios of RPKM values for the plus and minus genomic strands along the chromosome are shown for each pAgo.

To measure the effects of pAgos on the bacteriophage replication, we determined phage titers (the number of plaque forming units) at different times post infection (2 or 7 hours) in the control strain and in the strains with pAgos. It was found that DloAgo or CepAgo did not change the number of infectious phage particles, which correlated with the absence of their effects on cell growth during infection (Fig. 6B and C). EmaAgo did not strongly affect phage titers at 2 hours post infection but significantly decreased them at 7 hours post infection at all tested MOI (34-fold, 50-fold and 49-fold at MOI= 0.00016, 0.0008 and 0.004, respectively) (Fig. 6C).

We then measured the amount of phage DNA inside cells at different stages of phage infection in the control strain or in strains expressing EmaAgo or DloAgo. To analyze intracellular phage DNA content and remove free phage particles, the cells were extensively washed prior to DNA isolation. In the control strain and in the strain with DloAgo, the amount of phage DNA exponentially increased during infection. The ratio of phage to chromosomal DNA was increased from 0.001-0.004 at early stages of infection to >16 at 6 hours post infection (Fig. 7A). In contrast, in the strain expressing EmaAgo the amount of phage DNA was significantly lower than in the control strains starting from 3-4 hours post infection. Therefore, the observed decrease in the titer of infectious phage particles is explained by deficient phage replication during EmaAgo expression, likely as a result of phage DNA targeting by pAgo.

To test this hypothesis, we analyzed whether EmaAgo is loaded with smDNAs processed from the phage genome during infection. We purified EmaAgo from *E. coli* infected with P1, isolated and sequenced associated smDNAs, and mapped them to the *E. coli* and P1 genomes. It was found that phage-derived sequences are highly enriched in the smDNA library, when normalized by the relative length of these replicon and the copy number of phage DNA in the *E. coli* culture (Table S1). In particular, there are ~24-32 times more phage smDNAs than expected based on the measured phage to chromosomal DNA ratio (depending on the method used for calculation, Table S1). SmDNAs map to both strands of P1, with no preferences for specific genomic sites including origins of replication (*oriL* and *oriR*), *parS*, *loxP* and *pac* sites (Fig. 7B). This indicates intense processing of the whole phage genome into smDNAs loaded into EmaAgo.

**Figure 7.**
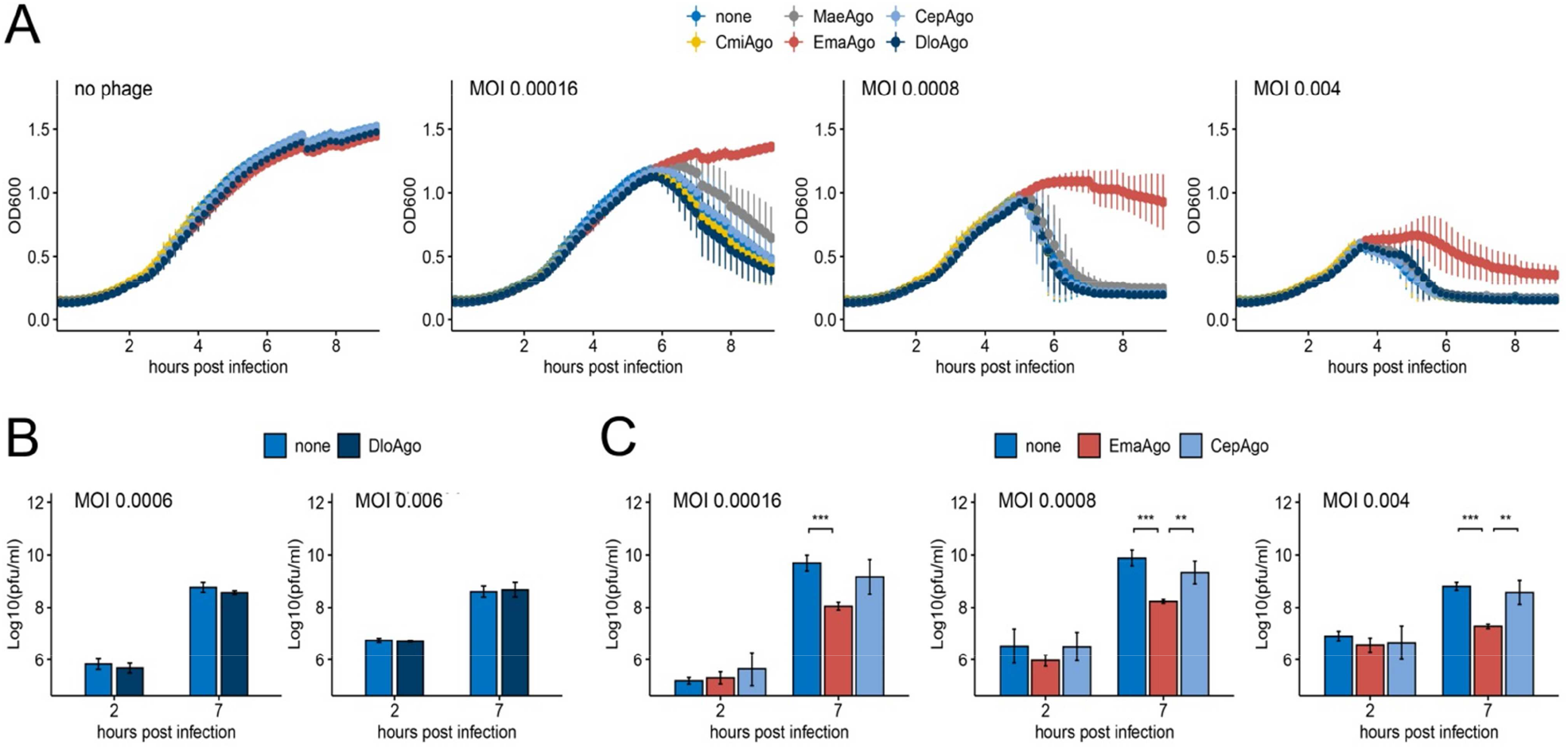
Effects of pAgos on infection of *E. coli* with phage P1. (A) Bacterial culture growth during P1 infection with different MOI in strains expressing plasmid-encoded pAgos and in the control strain without pAgos. Means and standard deviations from three independent experiments. (B) Titers of P1 at 2 or 7 hours after infection at different MOI (indicated in the figure) of the control strain or the strain expressing DloAgo. (C) The same experiment performed with strains expressing EmaAgo or CepAgo. PFU, plaque forming units. Means and standard deviations from three independent measurements (*p<0.05, ***p<0.001).

**Figure 8.**
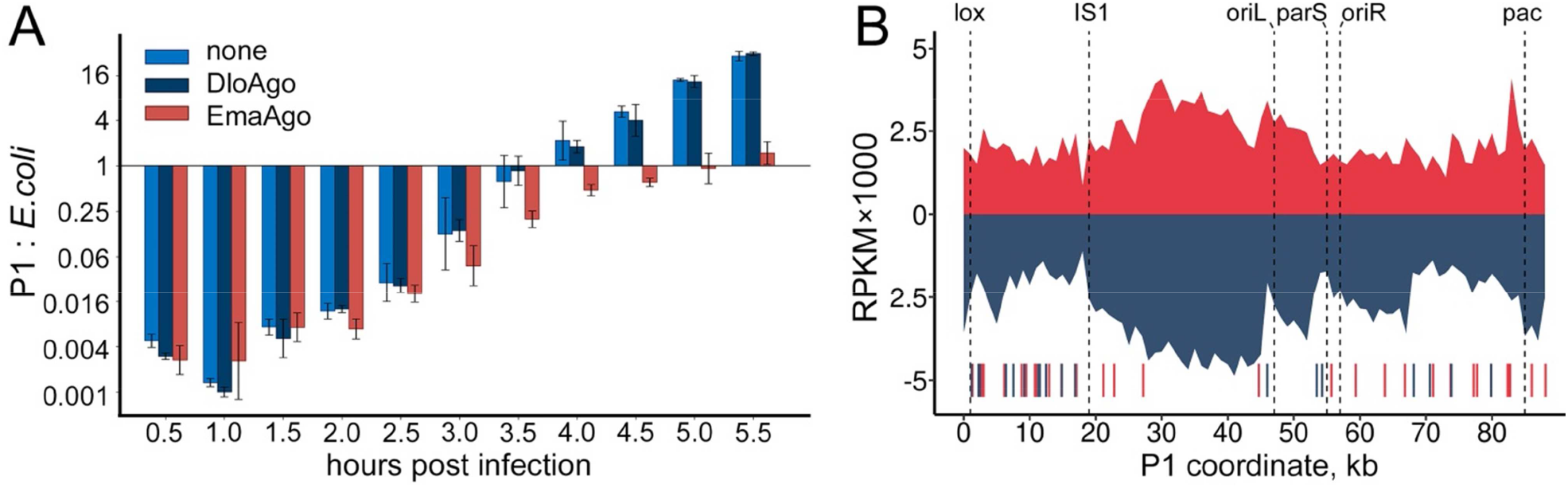
Targeting of phage DNA by EmaAgo. (A) Molar ratios of phage to chromosomal DNA in *E. coli* strains expressing DloAgo, EmaAgo, or lacking pAgos. Total DNA was isolated from the infected culture (MOI=7×10^−5^) at indicated times after infection and the amounts of DNA were measured by quantitative PCR for the phage and chromosomal loci as described in Materials and Methods. Means and standard deviations from three independent measurements. (B) Distribution of smDNAs along the P1 genome (red, plus strand; dark blue, minus strand). SmDNAs were isolated from EmaAgo at 2.5 hours post infection (MOI=0.0015) and sequenced. Positions of the phage origins of replications (oriL and oriR), the *pac*, *lox* and *parS* sites are indicated. SmDNA density is shown in RPKM values (reads per kilobase per million reads in the library) in 1 kilobase windows.

## Discussion

Analysis of pAgo proteins from diverse phylogenetic groups proved to be a fruitful approach to finding pAgos with various types of activities and allowed characterization of programmable nucleases with different specificities, unknown for eukaryotic Argonautes (see Introduction). In particular, previous studies identified several pAgo proteins from thermophilic and mesophilic prokaryotic species with DNA-guided DNA nuclease activities, which may have possible application in genome editing and biotechnology (Fig. 1A) (5,6,8-12,41). However, their functions in bacterial cells are not fully understood. It was recently shown that CbAgo from *C. butyricum* can protect the cells from phage infection (19), several pAgos affect plasmid transformation and maintenance (8-10,19), and TtAgo participates in DNA decatenation during replication (21), but whether these are universal activities for various groups of pAgo proteins have remained unknown. In this study, we have characterized five novel pAgo proteins from mesophilic bacteria which all are DNA-guided DNA endonucleases *in vitro* but have surprisingly different properties when expressed in bacterial cells *in vivo*.

*In vitro*, all five pAgos can precisely cut complementary DNA targets when programmed with small DNA guides in the range of 14-22 nucleotides and are active in a wide range of conditions, including physiological temperatures. Similarly to several previously studied pAgos (5,6,8,9,11,12), efficient target DNA cleavages requires the presence of the 5’-phosphate in guide DNA, but does not depend on the identity of the first nucleotide. DNA cleavage is also affected by mismatches around the cleavage site and in the 3’-supplementary region of guide DNA, similarly to other pAgos, including TtAgo, CbAgo, KmAgo and LrAgo (5,11,12,42). This contrasts eAgos, in which the seed region of guide RNA is most important for RNA target recognition (43), and may be a common feature of DNA-targeting pAgos. Furthermore, we have demonstrated that SSB can facilitate target DNA cleavage by some pAgos. A stimulatory effect of SSB on DNA cleavage was also previously reported for TtAgo (42), and SSB proteins were shown to be associated with SeAgo in the cells of *Synechococcus elongatus* (6). SSB may help to unfold the secondary structure of single-stranded DNA inhibitory for cleavage and can thus be generally used for increasing the cleavage efficiency of pAgo proteins *in vitro*. The analyzed pAgos are not very efficient in specific cleavage of double-stranded DNA, similarly to previously studied pAgos (5,9,32,33). In particular, EmaAgo preferentially performs nonspecific guide-independent cleavage of plasmid DNA *in vitro*. Such DNA chopping activity may potentially result in autonomous generation of large quantities of guide DNAs for further degradation of the target replicons.

*In vivo*, the five pAgo proteins reveal different patterns of DNA targeting and have different effects on phage infection in *E. coli*. At the chromosomal level, all five pAgos bind smDNAs corresponding to the *ter* sites of replication termination. Similar targeting of the chromosomal *ter* region was previously reported for CbAgo in *E. coli* cells, SeAgo in *S. elongatus* and TtAgo in *T. thermophilus* (6,19,21). TtAgo was shown to play a role in chromosome decatenation and completion of replication (21) but whether other pAgo proteins that also target the *ter* region may have similar functions remains to be established.

All analyzed pAgos are also loaded with smDNAs corresponding to the lambda prophage present in the expression strain, similarly to previously studied CbAgo and KmAgo (11,19). The DE3 prophage contains an inactivated integrase gene and is thought to be unable to excise its genome from the chromosome (44). However, the presence of a peak of pAgo-associated small DNAs centered at the left prophage end indicates genomic DNA processing in this region. Such processing might potentially result from partial prophage excision due to an unanticipated activity of the mutant integrase, which requires further investigation.

Unlike the other four pAgos, EmaAgo can induce DNA interference between homologous plasmid and chromosomal sequences, resulting in generation of a smDNA peak around the *araC* gene. This indicates that EmaAgo, loaded with plasmid-derived guides, can cleave the homologous chromosomal gene followed by its further processing. Accordingly, EmaAgo is more efficiently loaded with plasmid-derived guide DNAs *in vivo*. EmaAgo also targets other regions of the chromosome, possibly as a result of DNA interference between multicopy chromosomal sequences. Previously, a similar pattern of genomic DNA targeting was observed for CbAgo when it was expressed in *E. coli* (19).

Generation of guide DNAs associated with the analyzed pAgos to various degrees depends on the activity of the RecBCD helicase-nuclease. RecBCD likely plays the main role in biogenesis of smDNAs produced at *ter* sites, in accordance with its proposed functions in replication termination (35–37). RecBCD also makes an important contribution to generation of smDNAs around the *araC* gene during DNA interference in the case of EmaAgo and previously studied CbAgo (19). Furthermore, whole-genome analysis of smDNAs demonstrated that they are produced asymmetrically around Chi sites in the case of EmaAgo, DloAgo and CbAgo, indicating that RecBCD is also involved in the biogenesis of smDNAs from multiple chromosomal regions. In contrast, analysis of smDNAs associated with the remaining three pAgos, CmiAgo, MaeAgo and CepAgo, does not reveal Chi-dependent asymmetry. This suggests that pAgo proteins rely on the homologous recombination machinery for smDNA generation to various extent.

Intriguingly, Chi-dependent asymmetry in smDNA processing is observed only for one of the two DNA strands, co-oriented with the Chi sequence, for both individual peaks of smDNAs and metaplots generated from the whole-genome data. This indicates that the two DNA strands are asymmetrically cleaved by RecBCD and, possibly, other nucleases during double-strand break processing. One explanation may be that RecBCD degrades the 3’-terminated and 5’-terminated strands of a double-stranded end in different ways (38,40). This may result in preferential generation of smDNAs from the products of degradation of the 3’-terminated strand, which are then bound by pAgos, until processing stops at the Chi site. According to another model, RecBCD unwinds DNA without cleavage until the recognition of Chi, where it cuts the 5’-terminated strand (39). In this case, preferential generation of pAgo-bound smDNAs from the 3’-terminated strand may result from the action of an additional cellular nuclease(s), such as RecJ. In both cases, smDNAs corresponding to the 5’-terminated strand may be generated by other nucleases independently of Chi sites and RecBCD.

Previously, chromosomal DNA processing by two studied pAgo proteins, CbAgo and TtAgo, was proposed to be connected to DNA replication and repair. TtAgo promotes chromosome decatenation likely by introducing DNA breaks at the *ter* region (21). CbAgo was shown to be actively loaded with smDNAs from engineered double-strand breaks (19).

The new pAgos, including EmaAgo and DloAgo, also bind the products of double-strand break processing, as suggested by the patterns of smDNA loading around Chi sites. Furthermore, these pAgos reveal subtle but recognizable differences in DNA targeting during replication, with preferential loading of smDNA from either the lagging (EmaAgo and CepAgo) or leading (DloAgo) DNA strands. CbAgo preferentially binds smDNAs derived from the lagging genomic strand. This indicates that smDNA processing is dependent on replication and that pathways of such processing may be different for various pAgos. It can be proposed that the observed differences in bacterial DNA targeting by pAgos may result from their different interactions with the DNA substrates and/or from differences in their cooperation with cellular machineries involved in DNA replication, processing and repair.

EmaAgo, but not other pAgos analyzed in this study, protects bacterial cells from phage infection. Analysis of the phage DNA content demonstrated that EmaAgo does not have strong effects on phage replication at early stages of infection, before detectable cell lysis. However, it strongly decreases the amount of intracellular phage DNA and the number of infectious phage particles at later stages, suggesting that it may attack the phage replication intermediates during the active phase of infection. Phage-derived DNA fragments are strongly enriched among smDNAs bound to EmaAgo and cover the entire phage sequence demonstrating intense processing of the phage genome.

EmaAgo and other analyzed pAgos are also enriched with smDNAs generated from plasmid sequences, but the antiphage activity is observed only for EmaAgo. Importantly, neither of the tested pAgos except EmaAgo cleaves homologous loci in the process of DNA interference. At the same time, CbAgo was previously shown both to induce DNA interference between multicopy loci and to protect *E. coli* from invader DNA (19). Therefore, the ability of pAgos to defend cells from infection apparently correlates with their ability to induce DNA interference between multicopy DNA elements. RecBCD and homologous nucleases (such as AddAB in some species) may contribute to preferential processing of invader elements, because of their intense replication, small replicon sizes and a lack of Chi sites that protect chromosomal DNA by promoting repair of double-strand breaks by homologous recombination (19,45). Thus, initial non-sequence specific processing of foreign replicons by cellular nucleases can be converted into their specific targeting by pAgo nucleases by generating small DNA guides bound and used by pAgos. Potentially, similar guide-directed targeting of homologous sequences by pAgos may also stimulate DNA recombination and repair. Whether pAgos can contribute to these processes in various bacterial species remains to be studied.

## Data Availability

The smDNA sequencing datasets generated in this study are available from the Sequence Read Archive (SRA) database under accession numbers PRJNA827032 and PRJNA827167. The code used for data analysis is available at the GitHub repository at https://github.com/AlekseiAgapov/5pAgos. The genomic sequence of phage P1 used in the experiments is available from GenBank under accession number OP279344. All primary data are available from the corresponding author upon request.

## Funding

This study was supported in part by the Russian Science Foundation (grant 20-74-10127 to AA, analysis of *in vitro* activities of pAgos; grant 19-14-00359 to DE, analysis of genomic DNA targeting by pAgos).

## Acknowledgements

We thank Dr. David Leach and Dr. Gerry Smith for insightful discussions, Dr. Anna Olina for help in preparation of smDNA libaries.

## Conflict of interest

The authors declare no conflict of interests.

## Notes

### Competing Interest Statement

The authors have declared no competing interest.

