## Supplemental Info for "Bacterial Argonaute nucleases reveal different modes of DNA targeting *in vitro* and *in vivo*"

### **SUPPLEMENTARY INFORMATION**

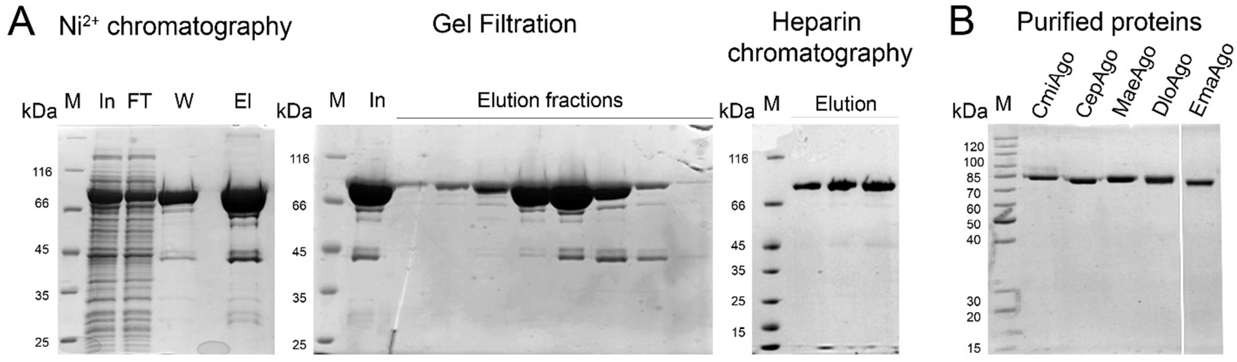

**Figure S1. Purification of pAgos.** (A) Illustration of individual purification steps for DloAgo. Input (In), flow-through (FT), wash (W) and elution (El) fractions are shown. (B) Electrophoretic analysis of the final pAgo preparations used in this study. The final protein samples were >98% pure based on Coumassie staining.

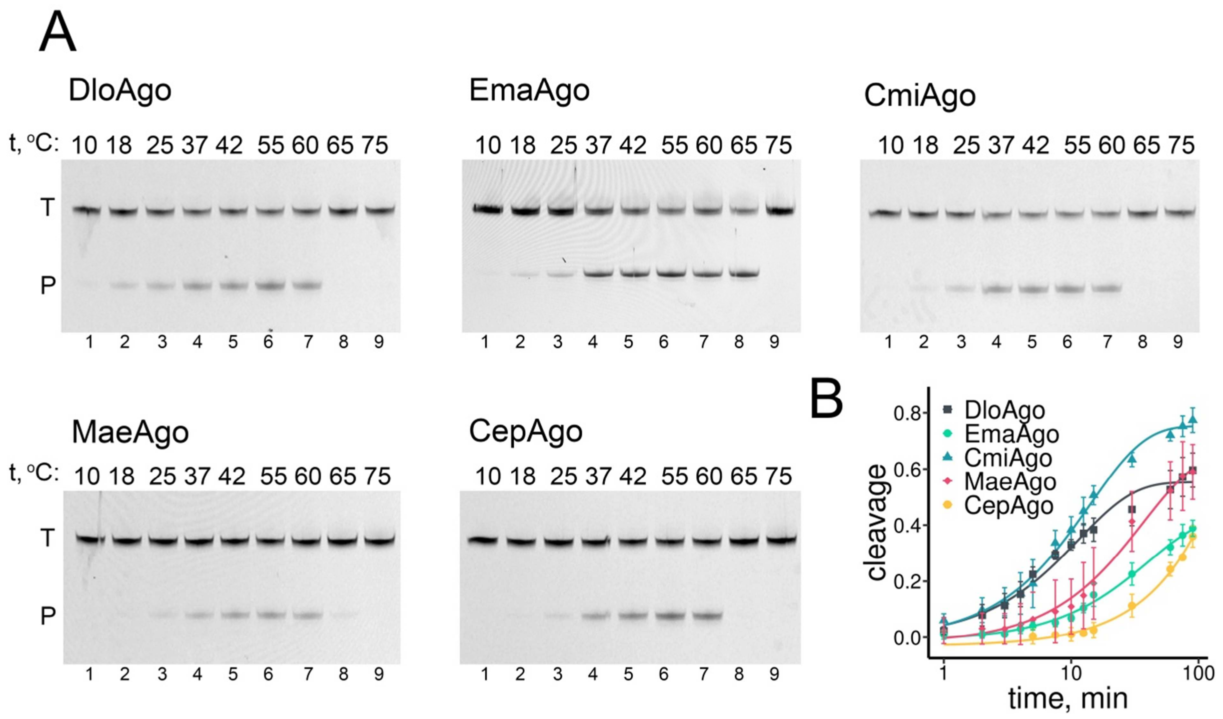

**Figure S2. Catalytic properties of the five pAgos proteins.** (A) Temperature dependence of the nuclease activity of various pAgos proteins. The reactions were performed at indicated temperatures using standard guide and target DNA (3'-Cy5-labeled) oligonucleotides (Table S2). Positions of the 3'-labeled target (T) and product (P) oligonucleotides are indicated. (B) Catalytic rates of different pAgos under single-round conditions (1 μM pAgo, 250 nM guide DNA, 50 nM target DNA). The reactions were performed with 3'-Cy5-labeled target DNA for indicated time intervals at 37 °C and the data were fitted to a single exponential equation. Means and standard deviations from three independent experiments are shown. The resulting $k_{obs}$ values are 0.085±0.016 $min^{-1}$ for DloAgo, 0.023±0.008 $min^{-1}$ for EmaAgo, 0.073±0.023 $min^{-1}$ for CmiAgo, 0.029±0.014 $min^{-1}$ for MaeAgo, 0.0045±0.0003 $min^{-1}$ for CepAgo.

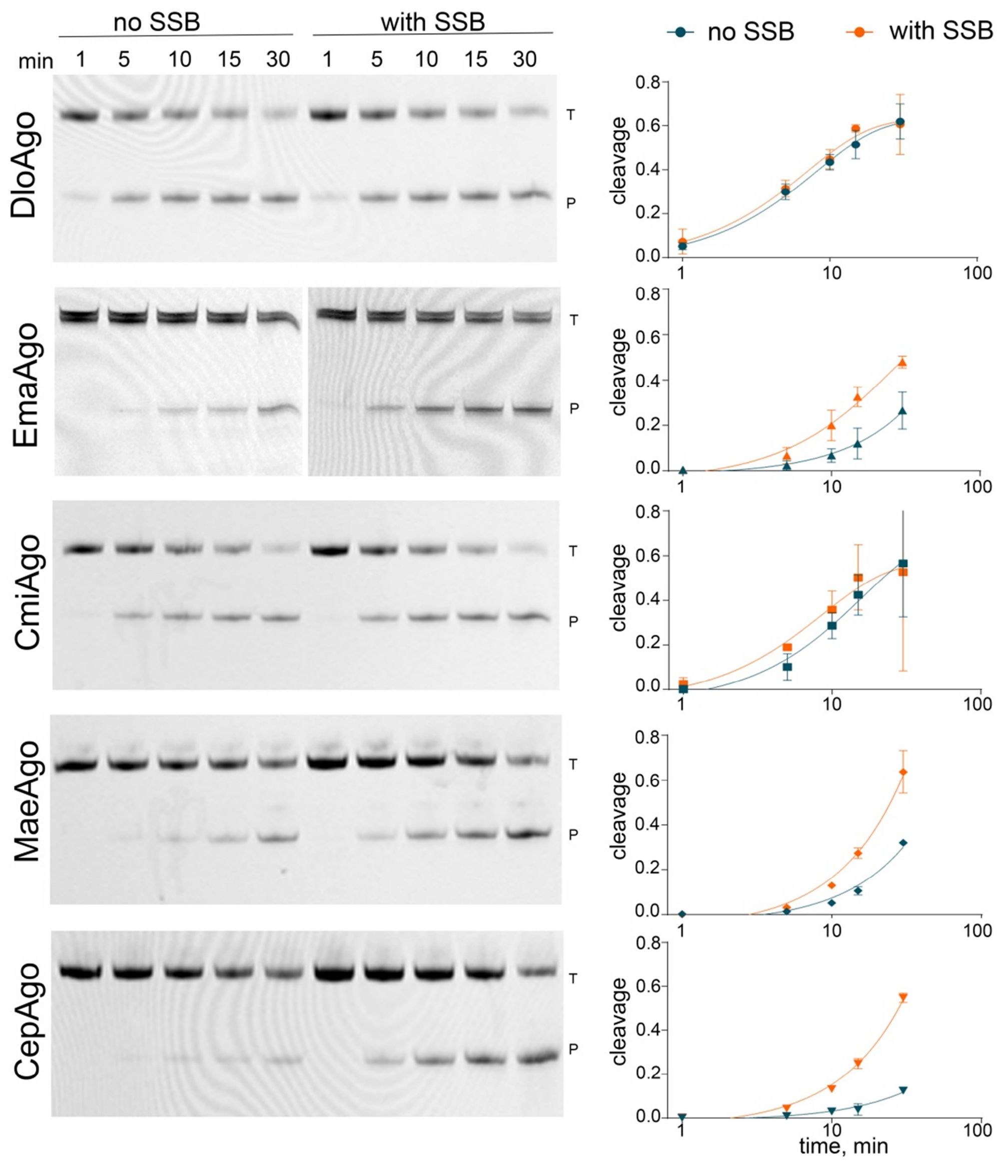

**Figure S3. Effects of *E. coli* SSB on target DNA cleavage by different pAgos.** (A) Analysis of the cleavage efficiencies after different incubation times using a 3'-Cy5-labeled target DNA. The reactions were performed with indicated pAgos at 37 °C in the absence or in the presence of *E. coli* SSB. Positions of the 3'-labeled target (T) and product (P) oligonucleotides are indicated. (B) Kinetics of DNA cleavage by various pAgos depending on the presence of SSB. Means and standard deviations from three independent experiments are shown.

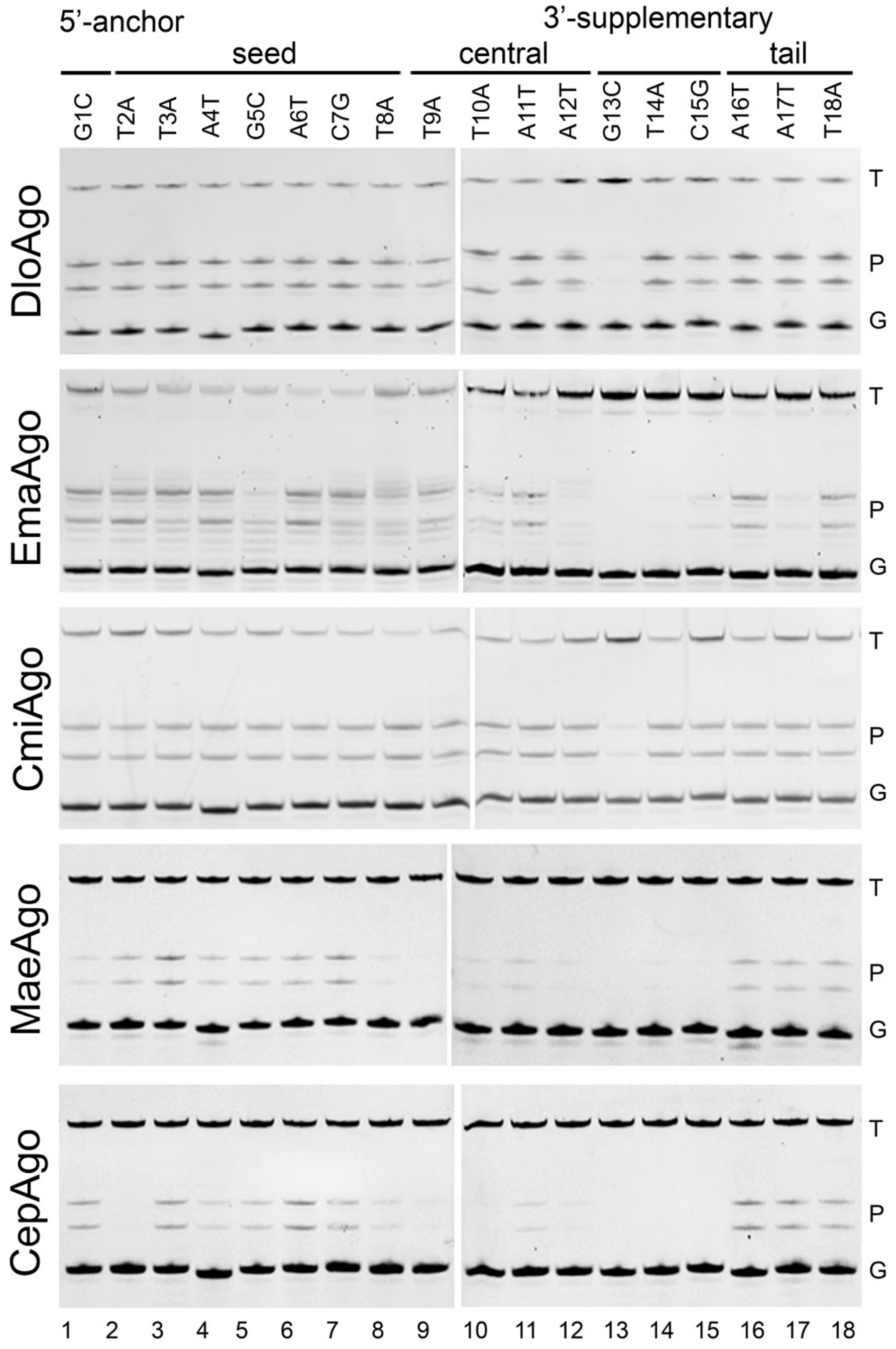

**Figure S4. Effects of mismatches between the guide and target DNA on the efficiency of DNA cleavage by pAgos.** DNA cleavage was analyzed using a set of guide DNAs containing mismatches with the same target at each individual position (substitutions in the guide oligonucleotide are indicated on the top; see Table S2 for oligonucleotide sequences). Representative gels from 1-3 replicate experiments are shown. Positions of the target (T), guide (G) and product (P) oligonucleotides are indicated.

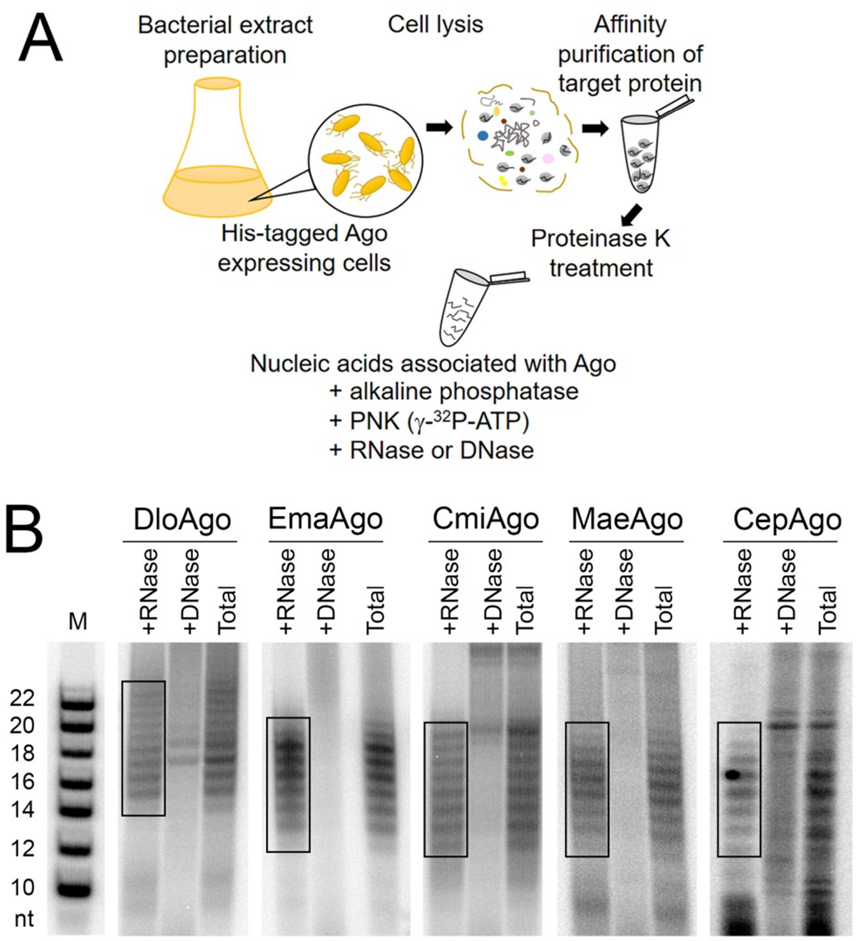

**Figure S5. Analysis of smDNAs associated with pAgos during their expression in *E. coli*.** (A) Scheme of the experiment. (B) Electrophoretic analysis of labeled smDNAs purified from pAgos (19% denaturing urea PAGE). Nucleic acids isolated from pAgos were treated with alkaline phosphatase to remove pre-existing 5'-phosphates and 5'-labeled with $\gamma$-$P^{32}$-ATP and polynucleotide kinase. After labeling, the samples were treated with either RNase A, or with DNase I, or left untreated ('Total' fraction). The marker lane (M) contains 5'-labeled DNA oligonucleotides of indicated lengths. For each pAgo, the range of smDNAs used for sequencing is indicated.

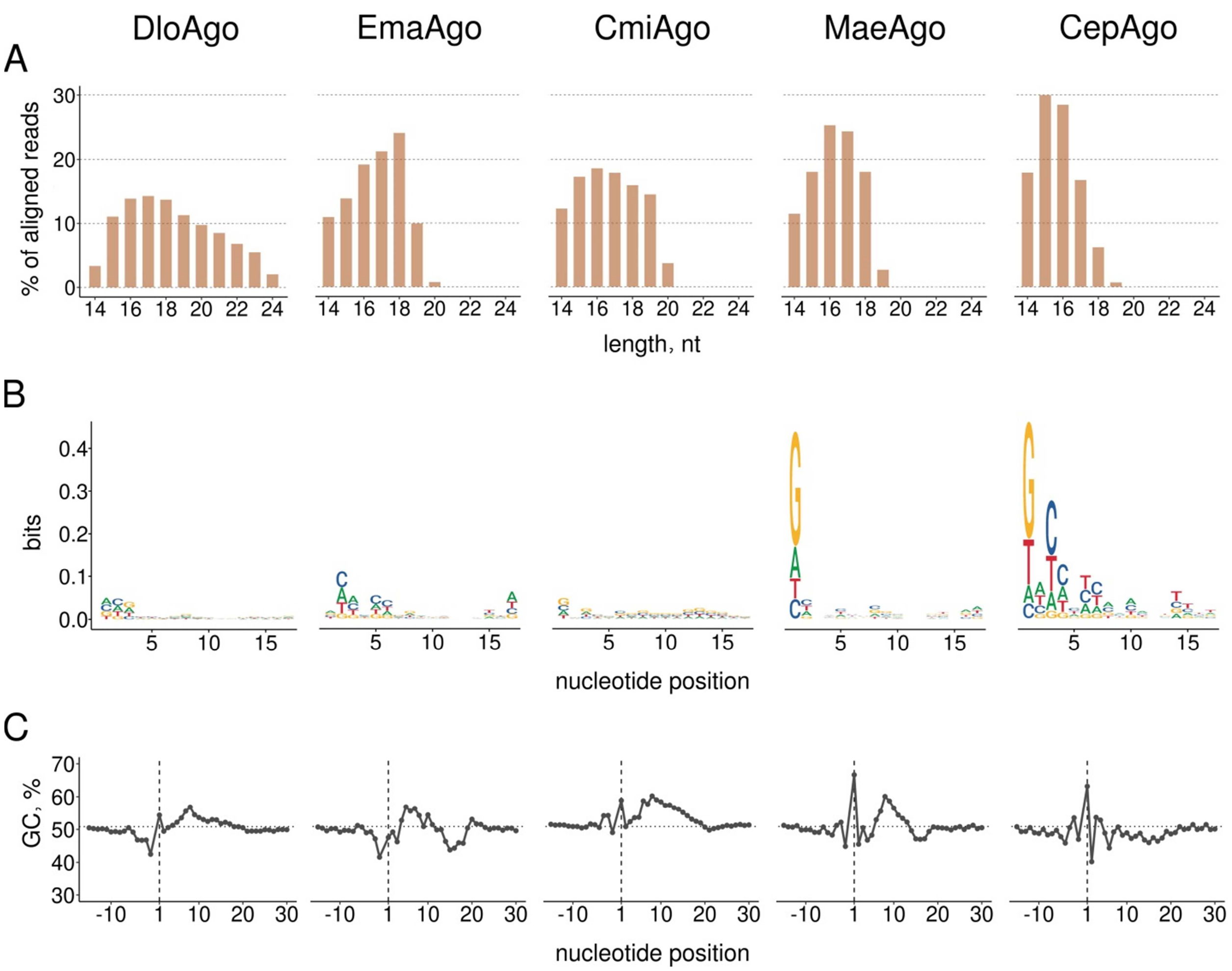

**Figure S6. Characteristics of smDNAs associated with pAgos.** (A) Length distribution in sequenced smDNA libraries for each pAgos protein. (B) Nucleotide biases along the smDNA sequences. (C) G/C-content in the smDNA sequences and in surrounding genomic regions.

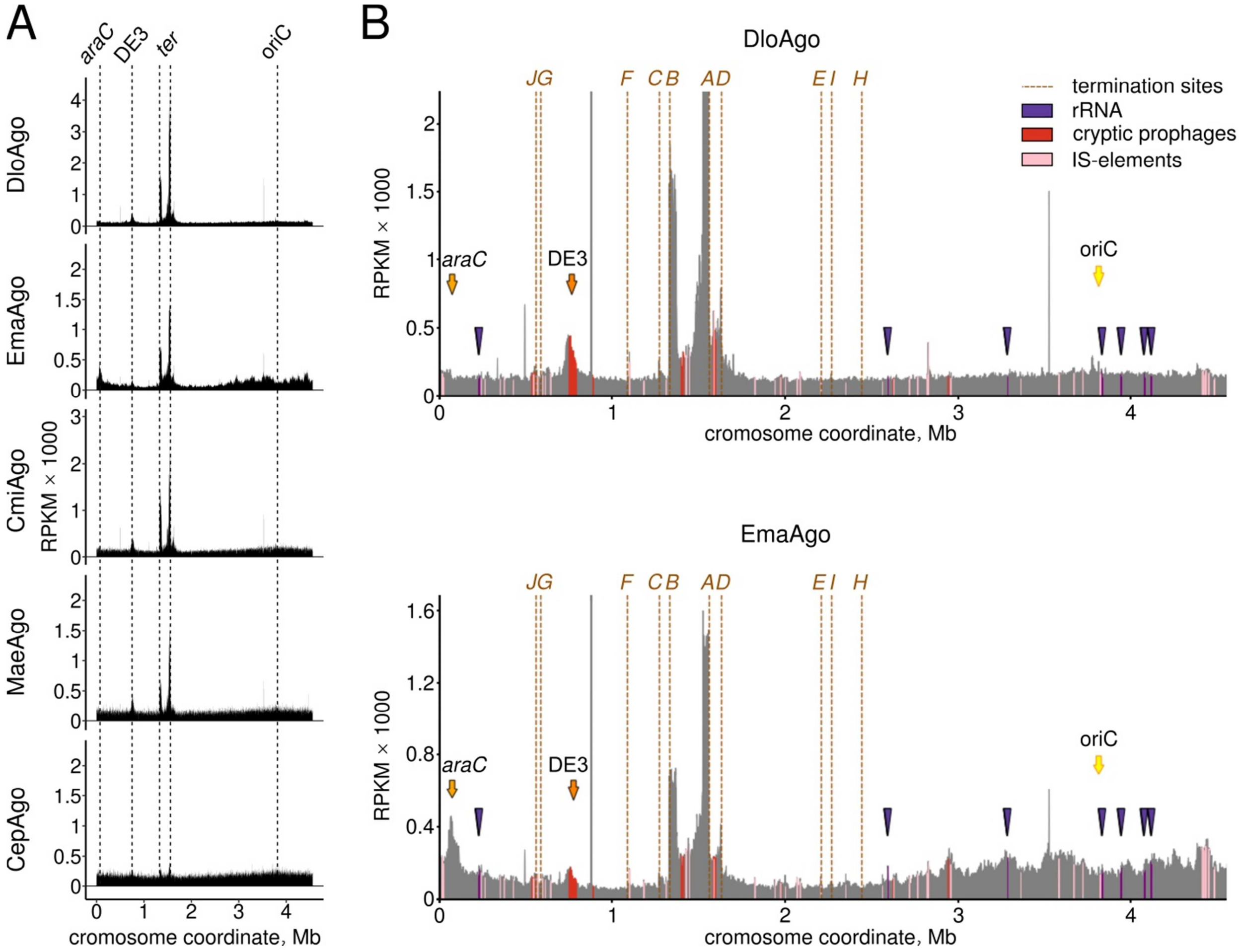

**Figure S7. Whole-genome distribution of smDNAs associated with pAgos.** (A) Distribution of smDNAs along the *E. coli* genome shown as the sum of reads corresponding to both genomic strands. Positions of *oriC*, *ter* region, *araC* and the DE3 prophage are indicated. (B) Enlarged views of the genomic distribution of smDNAs for DloAgo (top) and EmaAgo (bottom), showing positions of various genomic elements of *E. coli*: *ter* sites, dotted orange lines with uppercase letters; rRNA operons, violet triangles; prophages, red; IS-elements, pink.

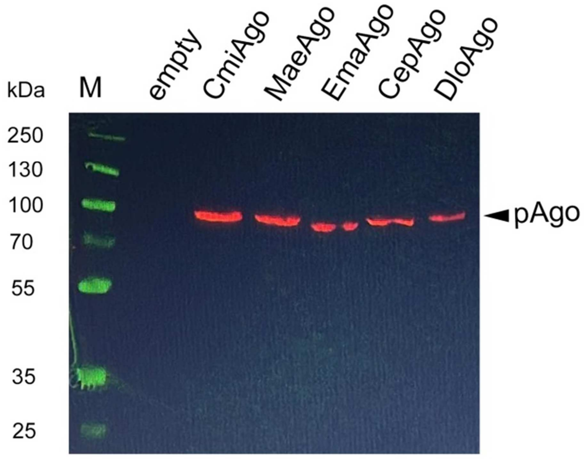

**Figure S8. Analysis of the expression of pAgo proteins in *E. coli* cells by Western blotting.** Ten microliters of night culture of MG1655 Z1 transformed with empty pBAD or pBAD encoding CmiAgo, MaeAgo, EmaAgo, CepAgo or DloAgo were inoculated in 1 ml of LB supplemented with 100 µg/ml ampicillin and 0.05% arabinose in a 12-well plate. The plate was incubated with shaking at 30°C for 2.5 hours and 500 µl of suspension was taken. The cells were precipitated and resuspended in 70 µl of $1^x$ Laemmli buffer and lysed at 95 °C for 5 min. Eight microliters of each sample were separated by 10% SDS-PAGE. The proteins were transferred to a nitrocellulose membrane and the membrane was blocked with 5% milk for 1 hour in the PBS-T buffer. The membrane was further incubated in the PBS-T buffer with 2.5% milk and mouse anti-His-tag antibodies (Sigma) overnight at 4 °C, washed twice with PBS-T and incubated with PBS-T supplemented with anti-mouse HRP-conjugate antibodies (Sigma) for 1 hour. The membrane was washed with PBS-T and the bands were visualized using the Clarity Max Western ECL substrate (Thermofisher) by a GelDoc documentation system (BioRad) in two channels, chemiluminescence and colorimetric (for visualization of molecular weight markers).

**Table S1. Analysis of the enrichment of pAgo proteins with plasmid and phage smDNAs.** Small DNAs were isolated from pAgos purified from *E. coli* BL21(DE3) expressing each pAgo at 4 hours after induction of pAgo expression or from *E. coli* MG1655 expressing EmaAgo at 2.5 hours after infection with phage P1vir (MOI=0.0015) and used for sequencing. The amounts of smDNAs mapped to the phage and chromosomal sequences were normalized by the relative lengths of the *E. coli* and P1 genomes and by the measured copy number of P1. Total genomic DNA was isolated from the same *E. coli* MG1655 culture and used for qPCR analysis or whole-genome sequencing. Expected number of reads = (*E. coli* genome mapped reads + plasmid/phage mapped reads)×(plasmid/phage size × copy number)/(plasmid/phage size × copy number + *E.coli* genome size)

| | |
|---|---|
| **DloAgo in *E. coli* BL21(DE3)** | |
| *E. coli* genome size, bp | 4558953 |
| smDNA reads mapped to the *E. coli* genome | 1755644 |
| **pBAD plasmid** | |
| plasmid size | 6315 |
| plasmid copy number | 12 |
| reads mapped to the plasmid | 274595 |
| expected number of plasmid reads | 33195 |
| **real/expected ratio for plasmid** | **8.3** |
| **EmaAgo in *E. coli* BL21(DE3)** | |
| *E. coli* genome size, bp | 4558953 |
| smDNA reads mapped to the *E. coli* genome | 2375747 |
| **pBAD plasmid** | |
| plasmid size | 6315 |
| plasmid copy number | 12 |
| reads mapped to the plasmid | 1139950 |
| expected number of plasmid reads | 56570 |
| **real/expected ratio for plasmid** | **20.2** |
| **CmiAgo in *E. coli* BL21(DE3)** | |
| *E. coli* genome size, bp | 4558953 |
| smDNA reads mapped to the *E. coli* genome | 612296 |
| **pBAD plasmid** | |
| plasmid size | 6315 |
| plasmid copy number | 12 |
| reads mapped to the plasmid | 74575 |
| expected number of plasmid reads | 11234 |
| **real/expected ratio for plasmid** | **6.6** |
| **MaeAgo in *E. coli* BL21(DE3)** | |
| *E. coli* genome size, bp | 4558953 |

| | |
|---|---|
| smDNA reads mapped to the *E. coli* genome | 169921 |
| **pBAD plasmid** | |
| plasmid size | 6315 |
| plasmid copy number | 12 |
| reads mapped to the plasmid | 33760 |
| expected number of plasmid reads | 3338 |
| **real/expected ratio for plasmid** | **10.1** |
| **CepAgo in *E. coli* BL21(DE3)** | |
| *E. coli* genome size, bp | 4558953 |
| smDNA reads mapped to the *E. coli* genome | 1228695 |
| **pBAD plasmid** | |
| plasmid size | 6315 |
| plasmid copy number | 12 |
| reads mapped to the plasmid | 168177 |
| expected number of plasmid reads | 22793 |
| **real/expected ratio for plasmid** | **7.4** |
| **EmaAgo in *E. coli* MG1655 during P1 infection** | |
| *E. coli* genome size, bp | 4641652 |
| smDNA reads mapped to the *E. coli* genome | 3399303 |
| **pBAD plasmid** | |
| plasmid size | 6213 |
| plasmid copy number | 12 |
| reads mapped to the plasmid | 152857 |
| expected number of plasmid reads | 56154 |
| **real/expected ratio for plasmid** | **2.7** |
| **Phage P1** | |
| phage size, bp | 88362 |
| reads mapped to phage DNA | 542583 |
| phage copy number (qPCR) | 0.31 |
| expected number of phage reads (qPCR) | 22955 |
| **real/expected ratio for phage (qPCR)** | **23.7** |
| phage copy number (shotgun) | 0.23 |
| expected number of phage reads (shotgun) | 17184 |
| **real/expected ratio for phage (shotgun)** | **31.6** |

**Table S2. Sequences of oligonucleotides used in the cleavage assays and in the quantitative analysis of phage DNA content.**

| Oligonucleotide | | Sequence (5'-3') | Description | Figure |
|---|---|---|---|---|
| **Cleavage assays** | | | | |
| 1. | G-guide-DNA | GTTAGACTTTAAGTCAAT | 18 nt guide DNA with 5'-G, complementary to G-target | 2B,C<br>3B |
| 2. | G-target-DNA | TTTATCAAAAAGAGTATTGACTT AAAGTCTAACCTATAGGATACTT ACAG | 50 nt target DNA for the guide/target specificity assay | 2B,C<br>3B<br>S4 |
| 3. | G-guide-RNA | GUUAGACUUUAAGUCAAU | 18 nt guide RNA for the guide/target specificity assay | 2B,C |
| 4. | G-target-RNA | UUUAUCAAAAAGAGUAUUGACU UAAAGUCUAACCUAUAGGAUAC UUACAG | 50 nt target RNA for the guide/target specificity assay | 2B,C |
| 5. | G-guide-10nt | GTTAGACTTT | 10 nt guide DNA with 5'-G, complementary to G-target | 3B |
| 6. | G-guide-12nt | GTTAGACTTTAA | 12 nt guide DNA with 5'-G, complementary to G-target | 3B |
| 7. | G-guide-14nt | GTTAGACTTTAAGT | 14 nt guide DNA with 5'-G, complementary to G-target | 3B |
| 8. | G-guide-16nt | GTTAGACTTTAAGTCA | 16 nt guide DNA with 5'-G, complementary to G-target | 3B |
| 9. | G-guide-20nt | GTTAGACTTTAAGTCAATAC | 20 nt guide DNA with 5'-G, complementary to G-target | 3B |
| 10. | G-guide-22nt | GTTAGACTTTAAGTCAATACTC | 22 nt guide DNA with 5'-G, complementary to G-target | 3B |
| 11. | +1NTguide(A) | AGAGCTGTCCCTCTCGAT | 18 nt guide DNA with 5'-A, complementary to Target_Kma (A) and Target_Kma (A)_cy5 | 3A<br>3C<br>S2A,B |
| 12. | +1NTguide(T) | TGAGCTGTCCCTCTCGAT | 18 nt guide DNA with 5'-T, complementary to Target_Kma (T) | 3C |
| 13. | +1NTguide(C) | CGAGCTGTCCCTCTCGAT | 18 nt guide DNA with 5'-C, complementary to Target_Kma (C) | 3C |
| 14. | +1NTguide(G) | GGAGCTGTCCCTCTCGAT | 18 nt guide DNA with 5'-G, complementary to Target_Kma (G) | 3C |
| 15. | Target_Kma (A)_Cy5 | AGGATACTTACAGCCATCGAGAG GGACAGCTC<u>T</u>ACTAGTCACCTGA GTCG-Cy5 | 3'-Cy5-labeled 50 nt target DNA for guide DNA with 5'-A, for assays with SSB, different cations, temperature, kinetics | 3A<br>S2A,B |
| 16. | Target_Kma (A) | AGGATACTTACAGCCATCGAGAG GGACAGCTC<u>T</u>ACTAGTCACCTGA GTCG | 50 nt target DNA for guide DNA with 5'-A | 3C |
| 17. | Target_Kma (T) | AGGATACTTACAGCCATCGAGAG GGACAGCTC<u>A</u>ACTAGTCACCTGA GTCG | 50 nt target DNA for guide DNA with 5'-T | 3C |
| 18. | Target_Kma (C) | AGGATACTTACAGCCATCGAGAG GGACAGCTC<u>G</u>ACTAGTCACCTGA GTCG | 50 nt target DNA for guide DNA with 5'-C | 3C |
| 19. | Target_Kma (G) | AGGATACTTACAGCCATCGAGAG GGACAGCTC<u>C</u>ACTAGTCACCTGA GTCG | 50 nt target DNA for guide DNA with 5'-G | 3C |

| | | | | |
|---|---|---|---|---|
| 20. | F1 guide | TTTTAAGTTGTTGATTTT | plasmid cleavage assay | 4C,D |
| 21. | R1 guide | TAAAAATCAACAACTTAA | plasmid cleavage assay | 4C,D |
| 22. | F2 guide | TAATAATTGACGATATGA | plasmid cleavage assay | 4C,D |
| 23. | R2 guide | GATCATATCGTCAATTAT | plasmid cleavage assay | 4C,D |
| **Mismatch assay** | | | | |
| 24. | G-guide_mm1 | CTTAGACTTTAAGTCAAT | guide forms mismatched pair in position 1 with G-target | S4 |
| 25. | G-guide_mm2 | GATAGACTTTAAGTCAAT | guide forms mismatched pair in position 2 with G-target | S4 |
| 26. | G-guide_mm3 | GTAAGACTTTAAGTCAAT | guide forms mismatched pair in position 3 with G-target | S4 |
| 27. | G-guide_mm4 | GTTTGACTTTAAGTCAAT | guide forms mismatched pair in position 4 with G-target | S4 |
| 28. | G-guide_mm5 | GTTACACTTTAAGTCAAT | guide forms mismatched pair in position 5 with G-target | S4 |
| 29. | G-guide_mm6 | GTTAGTCTTTAAGTCAAT | guide forms mismatched pair in position 6 with G-target | S4 |
| 30. | G-guide_mm7 | GTTAGAGTTTAAGTCAAT | guide forms mismatched pair in position 7 with G-target | S4 |
| 31. | G-guide_mm8 | GTTAGACATTAAGTCAAT | guide forms mismatched pair in position 8 with G-target | S4 |
| 32. | G-guide_mm9 | GTTAGACTATAAGTCAAT | guide forms mismatched pair in position 9 with G-target | S4 |
| 33. | G-guide_mm10 | GTTAGACTTAAAGTCAAT | guide forms mismatched pair in position 10 with G-target | S4 |
| 34. | G-guide_mm11 | GTTAGACTTTTAGTCAAT | guide forms mismatched pair in position 11 with G-target | S4 |
| 35. | G-guide_mm12 | GTTAGACTTTATGTCAAT | guide forms mismatched pair in position 12 with G-target | S4 |
| 36. | G-guide_mm13 | GTTAGACTTTAACTCAAT | guide forms mismatched pair in position 13 with G-target | S4 |
| 37. | G-guide_mm14 | GTTAGACTTTAAGACAAT | guide forms mismatched pair in position 14 with G-target | S4 |
| 38. | G-guide_mm15 | GTTAGACTTTAAGTGAAT | guide forms mismatched pair in position 15 with G-target | S4 |
| 39. | G-guide_mm16 | GTTAGACTTTAAGTCTAT | guide forms mismatched pair in position 16 with G-target | S4 |
| 40. | G-guide_mm17 | GTTAGACTTTAAGTCATT | guide forms mismatched pair in position 17 with G-target | S4 |
| 41. | G-guide_mm18 | GTTAGACTTTAAGTCAAA | guide forms mismatched pair in position 18 with G-target | S4 |
| **qPCR primers** | | | | |
| 42. | FE-primer | GGAACGCCATACCAGTCAGT | F-primer to *E. coli* genome | 8A |
| 43. | RE-primer | AAAGGCGCTAACTTCGACAA | R-primer to E. coli genome | 8A |
| 44. | FP1-primer | GATCCAGTTCCGTCAGCCAA | F-primer to phage P1 genome | 8A |
| 45. | RP1-primer | ATAACGCGGTAATCCGGTCC | R-primer to phage P1 genome | 8A |

**Table S3. Small DNA and genomic DNA libraries.**

| Library type | Strain | pAgo | Phage P1 | Growth Conditions | Growth Phase | Library type | BioProject accession number | SRA accession number |
|---|---|---|---|---|---|---|---|---|
| Small DNAs | *E. coli* BL21(DE3) | DloAgo | - | LB+Glucose+Ampicillin, arabinose induction, 37˚C -> 30˚C | 4h after induction | smDNA, SE50 | PRJNA827032 | SRR18769176 |
| | *E. coli* BL21(DE3) | EmaAgo | - | LB+Glucose+Ampicillin, arabinose induction, 37˚C -> 30˚C | 4h after induction | smDNA, SE50 | PRJNA827032 | SRR18771239 |
| | *E. coli* BL21(DE3) | CmiAgo | - | LB+Glucose+Ampicillin, arabinose induction, 37˚C -> 30˚C | 4h after induction | smDNA, SE50 | PRJNA827032 | SRR18770849 |
| | *E. coli* BL21(DE3) | MaeAgo | - | LB+Glucose+Ampicillin, arabinose induction, 37˚C -> 30˚C | 4h after induction | smDNA, SE50 | PRJNA827032 | SRR18770935 |
| | *E. coli* BL21(DE3) | CepAgo | - | LB+Glucose+Ampicillin, arabinose induction, 37˚C -> 30˚C | 4h after induction | smDNA, SE50 | PRJNA827032 | SRR18769219 |
| | *E. coli* MG1655 Z1 | EmaAgo | - | LB+Glucose+Ampicillin+Arabinose 30˚C | 2.5 h after mock infection | smDNA, SE50 | PRJNA827032 | SRR21098317 |
| | *E. coli* MG1655 Z1 | EmaAgo | + | LB+Glucose+Ampicillin+Arabinose 30˚C – 1st replica | 2h after infection | smDNA, SE50 | PRJNA827032 | SRR21098318 |
| | *E. coli* MG1655 Z1 | EmaAgo | + | LB+Glucose+Ampicillin+Arabinose 30˚C – 2nd replica | 2.5 h after infection | smDNA, SE50 | PRJNA827032 | SRR21177526 |
| Genomic DNA | *E. coli* MG1655 Z1 | no Ago | + | LB+Glucose+Ampicillin+Arabinose 30˚C | 2.5 h after infection | gDNA, SE200 | PRJNA827167 | SRR21091449 |
| | *E. coli* MG1655 Z1 | EmaAgo | + | LB+Glucose+Ampicillin+Arabinose 30˚C | 2.5 h after infection | gDNA, SE200 | PRJNA827167 | SRR21091647 |
